# Deep mutational scan of the pore of the cold-sensing TRPM8 channel

**DOI:** 10.64898/2026.04.28.721489

**Authors:** Averi Pike, Vincent Le, Ashlin Turner, Catalina Galván, Ashley Vazhavilla, Richard Salinas, Matt Bramble, Dennis C. Wylie, Jessica W. Podnar, Ian Hoskins, Can Cenik, Andrés Jara-Oseguera

**Author notes:** Correspondence should be addressed to: Andrés Jara-Oseguera. equal contributors (Averi Pike, Vincent Le, Ashlin Turner).

## Abstract

Members of the Transient Receptor Potential (TRP) family of ion channels have a nearly ubiquitous role in human physiology, tuning cell signaling to remarkably diverse physical and chemical stimuli. Although there is extensive structural data on TRP channels, a systematic and unbiased interrogation of structure-function relations in these proteins is required to fully elucidate their mechanisms of function. By focusing on a critical pore region of the TRPM8 channel, which is the main detector of cold and cooling agents in sensory neurons, we show how deep mutational scanning can be used in combination with the available structural data to understand how TRP channels respond to stimuli. We define a novel mechanism whereby the extracellular pore loop, which has only been resolved in structures representing desensitized states of the channel, plays an essential role in the response of TRPM8 to menthol or cold by coordinating the movement of the S6 helices that line and gate the pore, and the ion-selectivity filter that binds permeant cations. Moreover, our screen reveals sequence determinants along the S6 helices that explain how their architecture sustains gating and, together, provide strong support for a structural mechanism of TRPM8 pore opening in response to menthol and cold.

## Introduction

Mammals express 28 Transient Receptor Potential (TRP) ion channel genes across tissues and cell types^1^. Assembled mainly as homotetrameric complexes with a central ion-conducting pore^2^, TRP channels influence cellular signaling by enabling the movement of specific cations across membranes in response to diverse environmental signals. The TRPM8 channel, for example, functions as the main sensory detector of environmental cold in neurons and is also activated by cooling agents and natural products like menthol^3–5^. Like several other TRP channels^6^, TRPM8 is considered a polymodal receptor because its activity can be modulated by multiple types of stimuli including voltage^7,8^, phosphoinositide^9^ and other lipids^10^, G proteins^11,12^, and Ca^2+^ ions^13–15^, which interact with distinct regions within the receptor to modify its activity. Consequently, TRPM8 channel activity is highly dependent on the physiological and cellular context, which is important for processes like sensory adaptation in neurons^16^ or for the ability of TRPM8 to carry out functions in non-neuronal tissues where it is also expressed^17–20^. Conversely, TRPM8 dysregulation under pathological conditions can give rise to cold hyperalgesia^21^, migraine^22,23^, dry eye^24^, and even influence the aggressiveness of certain types of cancer^25–27^.

Almost three decades after TRPM8 was first cloned^5,28^, and despite an impressive amount of high-resolution structural data that has become available over the last ten years^14,15,29–33^, our understanding of the molecular basis for the polymodal nature of TRPM8 and other TRP channels, as well as their ability to respond to changes in temperature, remains incomplete. On the one hand, single-particle cryo-electron microscopy (cryo-EM) is limited in its ability to resolve dynamic protein regions as well as higher-energy states that may be essential components of the channel activation mechanism, while being susceptible to identifying states lacking biological significance. On the other hand, conventional approaches like site-directed mutagenesis combined with electrophysiology have insufficient throughput to interrogate the structural data in terms of a comprehensive mechanism of TRP channel function that considers the entire protein complex. Overcoming these challenges is critical to understanding TRP channels at the molecular level, because their polymodal nature necessitates contributions from multiple channel regions distributed across the whole protein to coordinate its responses to its diverse activators and regulators^34,35^. Notably, TRP channels are widely believed to detect heat or cold through multiple temperature-sensitive loci distributed across receptor complex^36^.

Deep mutational scanning approaches^37–43^ offer a promising emerging way to investigate structure-function relations in proteins at the scale that is required for large polymodal receptors like TRP channels. We have therefore implemented a deep mutational scanning approach to screen the response to menthol and cold of all 912 single-residue variants localized within a critical region of the pore of the rat TRPM8 channel **(Fig. 1A, Extended Data Fig. 1A, B)**. Our screen provided extensive insight into a highly conserved region of TRPM8 that is critical for its function: the pore-lining S6 helices that are responsible for opening and closing the pore, and the extracellular pore region that connects the S6 helices with the ion-selectivity filter which coordinates permeant ions. Although this extracellular pore loop is known to be essential for TRPM8 function^44^, its role in channel activation has remained unknown and its structure has only been resolved in structures of the channel in non-conducting states that are likely desensitized^14,15,29–32^. The deep mutational scanning results we obtained suggest a mechanism for how the extracellular pore loop contributes to function. The experimentally derived variant scores and the existing structural data^14,15,29–33^ together strongly support the described mechanism of pore opening in response to both cold and menthol, and reveals numerous sequence determinants that are fundamental to the dynamic character of the S6 helices and pore of TRPM8. Finally, we propose a mechanism of cold sensing involving the S6 helices in the pore.

**Figure 1.**
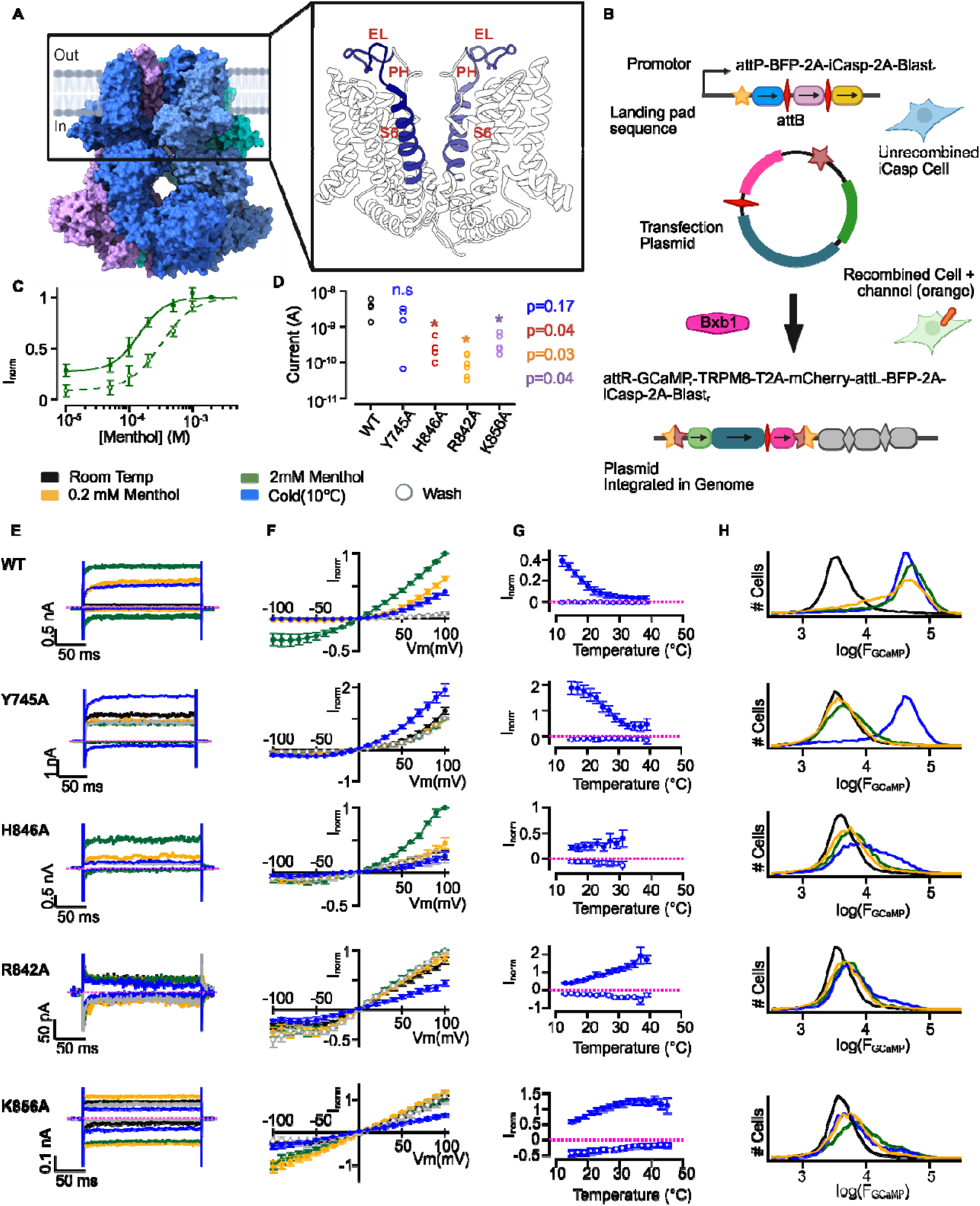
A Ca^2+^-fluorescence assay of TRPM8 channel activity. **(A)** Structural model of TRPM8 channel (left, PDB: 9B6D^15^) and the transmembrane domain (right) with the region between S927 and L974 shown in dark blue. EL, extracellular pore loop; PH, pore helix. **(B)** Cartoon of the iCasp9 landing pad expression system^46^, showing the genomic landing pad genes before recombination (top; star, attB sequence), the plasmid coding sequences (middle; star, attP recombination sequence; green, GCaMP6s; gray, IRES2; TRPM8, blue; red diamond, T2A; scarlet, mCherry), and the plasmid genes after Bxb1 (magenta shape)-dependent recombination (bottom). **(C)** Menthol concentration-response relation of TRPM8, measured at +100 (closed symbols) or -100 (open symbols) mV in whole cells. Data shown as mean ± SEM (n = 6). Curves are fits to the Hill equation (100 mV, EC_50_ = 0.15 ± 0.05 mM, Hill coef. = 1.50 ± 0.18; -100 mV, EC_50_ = 0.32 ± 0.06 mM, Hill coef. = 1.79 ± 0.09). **(D)** Current magnitude at 100 mV recorded at 10 °C (Y745) or at room temperature + 2 mM menthol (all other constructs) from individual cells expressing WT or mutant TRPM8. p-values shown are from a two-sided Student’s t-test between WT and each of the mutants. **(E)** Representative current traces from WT and mutant channels at ±100 mV recorded at each of the conditions denoted by the trace color. Dotted magenta lines denote the zero-current level. **(F)** Current voltage-relations of WT and mutants under the same conditions as in (E). Data shown as mean ± SEM (n = 4). **(G)** Current-temperature relations of WT and mutant TRPM8 measured at +100 (closed symbols) or -100 mV (open symbols). Data shown as mean ± SEM (n = 4). Dotted magenta lines denote the zero-current level. **(H)** GCaMP6s fluorescence intensity histograms of 10 x 10^3^ mCherry^+^ cells expressing WT or mutant channels measured by flow cytometry at each of the conditions denoted by the trace colors.

## Results

### A high-throughput fluorescence assay of TRPM8 response to cold and menthol

To carry out a deep mutational scan of the pore of TRPM8, we first set out to develop a high-throughput assay of channel activity. When expressed in the plasma membrane, TRPM8 activation leads to calcium entry into cells, which can be detected by an increase in the fluorescence intensity of a cytosolic Ca^2+^ reporter. We therefore co-expressed the rat TRPM8 channel together with the genetically encoded fluorescent Ca^2+^ reporter GCaMP6s^45^ and mCherry in iCasp9 landing pad cells^46^. This cell line enables stable expression of recombinant genes after integration of a promoter-less plasmid into a singular genomic landing pad^46^ **(Fig. 1B)**.

Using whole-cell patch clamp, we established that our wild type (WT) rat TRPM8 construct expressed in iCasp9 cells displayed menthol-, voltage-, and cold-dependent activation that was consistent with previous studies^5,47,48^ **(Fig. 1C-G)**. When analyzed by flow cytometry, we observed that exposure to cold or menthol of cells expressing WT TRPM8 channels caused a robust increase in GCaMP6s fluorescence **(Fig. 1H, top)**. To benchmark the sensitivity of this assay for detecting alterations in TRPM8 function, we examined four mutants with previously described phenotypes – Y745A that specifically disrupts menthol binding^49^, H846A and R842A that lower activation by both cold and menthol^50^, and the gain of function (GOF) K856A that has increased basal activity^50^. We first confirmed using patch clamp that the phenotype of each mutant agreed with the previous reports and found that R842A lacked channel activity altogether while H846A retained partial activity when fully stimulated **(Fig. 1E-G)**. When analyzed using the fluorescence assay, we confirmed that Y745A, H846A, and R842A had decreased GCaMP6s responses to menthol, and only Y745A responded normally to cold **(Fig. 1H)**, indicating that our assay successfully identified the negative alterations in function in each of these mutants. Notably, the assay did not detect major differences between R842A, H846A, and K856A. This was likely caused by the decreased cell surface expression of H846A and K856A, evidenced by their maximal currents that were significantly smaller than WT **(Fig. 1D)**. Therefore, we concluded that our Ca^2+^-fluorescence assay can be used to identify cells expressing TRPM8 variants having decreased activity in response to cold or menthol or decreased cell surface expression.

### Deep mutational scanning scores provide accurate insight into TRPM8 variant behavior

We focused our screen of the rat TRPM8 on a 48 amino acid region encompassing positions S927 to L974 **(Fig. 1A)**. This region is highly significant because it includes the poorly understood, yet functionally critical extracellular pore loop, and the S6 helices that form the walls of the pore and are responsible for its gating. We generated an expression plasmid library of all possible 912 single-reside mutants in this region using a method that allows nearly equal representation across variants^51^ **(Extended Data Fig. 1A, C)**, with 92% of variants within 2-fold of median depth. We then expressed this library in iCasp9 landing pad cells^46^ together with cells expressing four different synonymous variants that behaved like WT TRPM8 **(Extended Data Fig. 1B, H)**. The GCaMP6s fluorescence response of the cell library to menthol or cold was biphasic, indicative of altered activity or cell surface expression in a substantial portion of the variants **(Extended Data Fig. 1E)**. By flow cytometry we sorted single-, live-, and mCherry^+^-library cells into groups with low, medium, or high GCaMP6s fluorescence after exposure to cold or menthol **(Extended Data Fig. 1B, D-E)**.

In two independent biological replicates, we obtained enriched DNA amplicons of the channel coding region from each group of sorted cells and characterized the sequences by short-read NGS **(Extended Data Table 1)**. We quantified sequence reads coding for each of the synonymous and non-synonymous variants in the library^52^, and scored each variant using a Bayesian statistical method that assigns a numeric score and standard error to each variant based on its NGS read count frequency distribution across the low, medium, or high GCaMP6s-fluorescence groups^53^. In this scoring system, negative scores denote a reduction in channel expression or activity and scores closer to zero denote similarity to WT. The distribution of scores for both menthol and cold showed two components – a population of variants with the most negative scores, which were more narrowly distributed, while the rest of the variants had a wider score distribution **(Extended Data Fig. 1F)**. Importantly, the correlation between replicates of the scores for each variant was consistent with what has been obtained in other studies using similar methods^40,41,54^ **(Extended Data Fig. 1G)**. The NGS variant read counts that we used to calculate these scores were similarly distributed across groups of low, medium, or high GCaMP6s intensity in all experiments **(Extended Data Fig. 2A)**, and we did not observe a correlation between the total read counts for a variant and its score **(Extended Data Fig. 2B)**. Importantly, the total read counts per variant **(Extended Data Fig. 2C)** and the score standard errors **(Extended Data Fig. 2D)** were homogenous across most variants. These observations provide an initial technical validation to the reliability of the scores we obtained.

To benchmark the deep mutational scanning scores in terms of their cell surface expression or sensitivity to cold or menthol, we individually expressed and assayed 28 individual scored variants by their GCaMP6s fluorescence **(Extended Data Fig. 3A-C)**. We classified five variants as GOF because they had increased fluorescence at room temperature or in the presence of a half-activating concentration of menthol **(Extended Data Fig. 3C, blue)**, and eight variants as WT-like **(Extended Data Fig. 3C, grey)**. We also found five variants that had weaker responses to cold and menthol (mild loss of function, LOF) **(Extended Data Fig. 3C, light red)**, and ten variants with a strongly reduced response (strong loss of function) **(Extended Data Fig. 3C, dark red)**. By comparing these phenotypes with the scores for each variant, we established the following criteria to interpret the results of our screen: (1) GOF variants cannot be identified by our screen, because their fluorescence response in the presence of cold or 2 mM menthol can only be as high as that of WT or lower due to reduced expression and cytotoxicity **(Extended Data Fig. 3C, blue)**; (2) Individual variants with a score less than -1.5 are more likely to have altered gating or reduced cell surface expression, because 18 out of 20 scores for the strong LOF variants were smaller than -1, and 15 out of 20 were smaller than -1.5 **(Extended Data Fig. 3C, dark red)**; (3) Positions in the primary sequence where most scores are less than -1.5 are most likely to include variants with altered function or expression. The residues at these positions are likely essential for function or expression; (4) Positions where most scores are larger than -1 are most likely to include WT-like variants, because all WT-like variants had scores larger than -1 **(Extended Data Fig. 3C, gray)**. The residues at these positions are unlikely to contribute to channel function or expression; (5) Positions in the primary sequence where scores are distributed across the entire range, as we observed for the mild LOF variants **(Extended Data Fig. 3C, light red)**, are likely to include variants with altered function or expression. The residues at these positions most likely play a role in channel function or expression. This wider distribution of variant effects may be due to more relaxed structural constraints required for function at these sites, or to the role of these sites tuning channel folding, or its response to stimuli.

### The pore of TRPM8 responds similarly to cold and menthol

A color heat map of the deep mutational scanning scores revealed clear patterns in the variant effects along the primary sequence of the extracellular pore loop and S6 helices of TRPM8, which were nearly indistinguishable between the cold and menthol datasets **(Fig. 2A)**. Consistently, the median cold or menthol scores per position were similar across the entire sequence **(Fig. 2B)**, and we observed a strong correlation between the menthol and cold scores **(Fig. 2C)** that was comparable to the correlation between replicate cold or menthol experiments **(Extended Data Fig. 1G)**. Together, these observations indicate that despite cold and menthol being such different types of stimuli, variants in this region of the channel had comparable responses to either stimulus, suggesting that the pore undergoes similar conformational changes when activated by cold or menthol.

**Figure 2.**
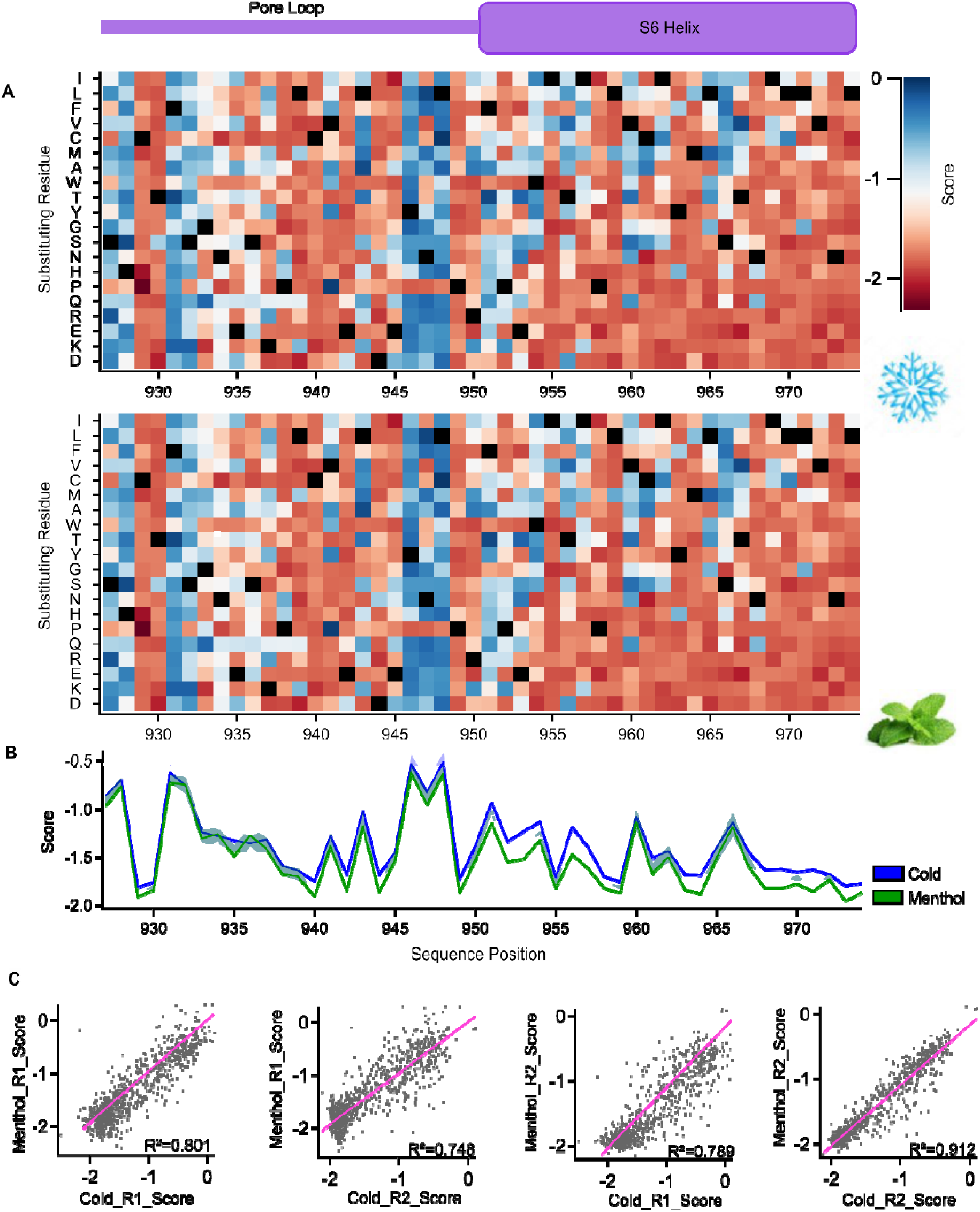
Deep mutational scanning scores of TRPM8 pore variants. **(A)** Heatmap of the individual variant scores obtained from cells stimulated with cold (top) or 2 mM menthol (bottom). Black squares denote WT residues. **(B)** Median score per sequence position for the cold and menthol screens. The shading is the SEM of the scores across the 19 variants per position. **(C)** Individual variant score correlations across experimental conditions and two biological replicate experiments.

### The essential role of the extracellular pore loop in TRPM8 channel gating

In the desensitized state of TRPM8^15^, the extracellular pore loop is shaped like the number eight and lies parallel to the plane of the membrane; the edge of the loop that is distal from the S6 helix forms an interface with the pore helix (P.H.) while the N- and C-terminal ends of the pore loop connect to the ion selectivity filter (S.F.) and S6 helix, respectively **(Fig. 3A)**. This fold is locked in place by a disulfide bond formed between C929 and C940 and shown to be essential for gating^44^. Notably, the pore loop and S6 helices had a nearly identical fold across 16 structures that were obtained under very different experimental conditions and that represent non-conducting, likely desensitized states of TRPM8 from mice, humans, and birds **(Extended Data Fig. 4A)**. To learn how the extracellular pore loop participates in TRPM8 function, we obtained cumulative histograms of all the scores at each position along the pore loop **(Fig. 3B)**, and used these to group positions into three categories: (1) Positions that are highly tolerant to substitution and that likely play no important role in channel function **(Fig. 3B, C; grey)**; (2) Critical positions where most or all substitutions strongly affected gating or expression **(Fig. 3B, C; dark red)**, which included C929 and C940; (3) Positions with a wide score distribution **(Fig. 3B, C; pink)**.

**Figure 3.**
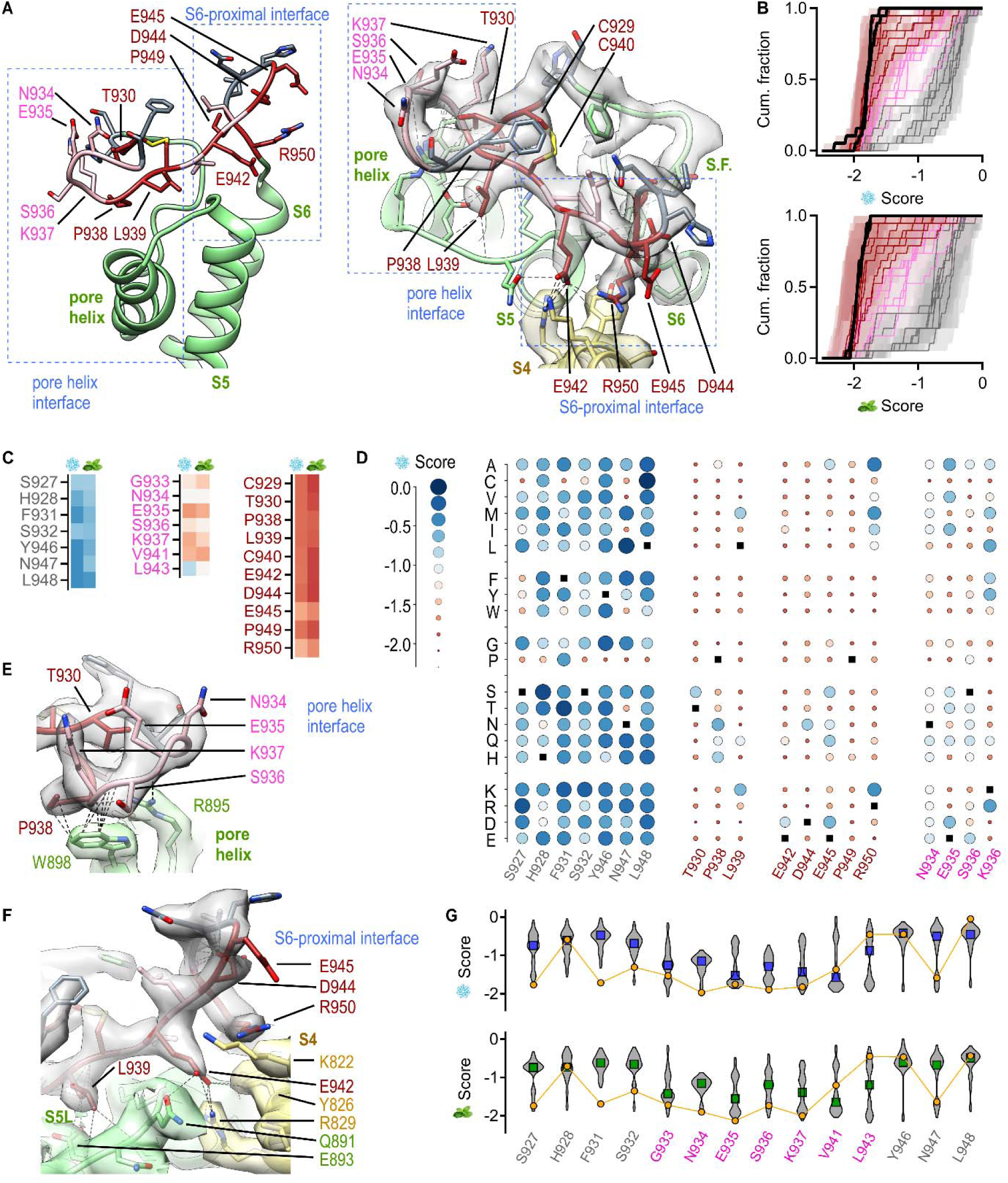
Sequence determinants in the extracellular pore loop of TRPM8. **(A, left)** Structural model of TRPM8 in the desensitized state^15^ (PDB: 9B6D) as seen from the side. Pore loop residues 927-950 are colored based on their low (dark red), intermediate (pink), or high tolerance of substitutions (gray). The pore helix and ion-selectivity filter (S.F.) of the same subunit as the pore loop are shown in green, and the S4 helix of the adjacent subunit in sand. **(A, right)** Same model as in the left panel seen from above, and including the electron microscopy (EM) map^15^ (EMD-44255). Dashed blue rectangles denote the pore helix and S6-proximal interaction interfaces between the pore helix and the rest of the protein. Dashed black lines in the right panel denote non-covalent interactions between the pore loop and the rest of the protein. **(B)** Cumulative histograms of all variant scores per sequence position, colored by their high (gray), intermediate (pink), or no tolerance (dark red) of substitutions. Shading is the average Lilace^53^ standard error per position. Histograms for C929 and C940 are shown as continuous black lines. **(C)** Variant classification and median score color scale as in Fig. 2A. **(D)** Bubble plot of individual variant cold scores. Scores are represented both by the radius of the circles and their color as denoted by the scale on the left and as displayed in Fig. 2A. Black squares denote WT residues. **(E)** Side view of the atomic model and EM map of the pore helix interface of the pore loop colored as in (A). **(F)** Side view of the atomic model and EM map of the S6-proximal pore loop interface with colors as in (A). Dashed black lines in (E) and (F) denote non-covalent interactions between the pore loop and the rest of the protein. **(G)** Violin plots of the individual variant scores at positions with intermediate and high tolerance of substitutions, relative to the individual score of the cysteine-containing variants at each position (yellow). Median scores shown in blue (cold) and green (menthol).

Upon mapping the three residue categories onto a desensitized state structure^15^ **(Fig. 3A)**, we noted that positions most tolerant of substitutions **(Fig. 3D, Extended Data Fig. 4B; gray)** were grouped together into three clusters in the structure with their sidechains facing the solvent. Based on the criteria we established for interpreting the scores, residues at these positions most likely do not contribute to channel function and retain a similar orientation towards the solvent during channel gating. Consistent with this conclusion, we note that the primary sequence of the pore loop is highly conserved between TRPM8 orthologues from humans to amphibians except at four short loci that include each of the three clusters mentioned above **(Extended Data Fig. 4C, D)**.

The residues that were most intolerant of substitutions that we deemed critical for channel function **(Fig. 3D, Extended Data Fig. 4B; dark red)** localized to two different interfaces between the pore loop and the rest of the protein: T930, P938, L939 on the S6-distal pore helix-interface **(Fig. 3A, E)**, and E942, D944, E945, P949, and R950 on the S6-proximal interface where the C-terminal end of the pore loop connects to the S6 helix **(Fig. 3A, F)**. The pore helix interface in the desensitized state is formed by interactions between residues in the pore helix and S5 helix linker and either the pore loop main chain or the non-polar side chains of P938 and L939, while interactions between T930 and the closely apposed pore loop main chain likely contribute to stabilizing the fold **(Fig. 3A, E)**. Due to the critical role of these residues based on our scores, it is likely that in addition to stabilizing the desensitized state, similar interactions at this pore helix interface are required to stabilize other functionally relevant states of the channel. This pore helix interface also includes four polar or charged residues, N934-K937, that face the solvent **(Fig. 3A, E)** and have an intermediate tolerance of substitutions **(Fig. 3D, Extended Data Fig. 4B; pink)**. The relation between the type of amino acid substituent and the resulting variant scores was starkly different at each of these four positions **(Fig. 3D, Extended Data Fig. 4B)**, indicating that each of these sites experiences distinct structural constraints during channel gating. We speculate that although not entirely essential, these residues may have an important role tuning the response of TRPM8 to cold and menthol. Consistent with this interpretation, glycosylation at position N934 has been shown to modulate the sensitivity of TRPM8 to cold and menthol^55^.

The S6-proximal interface at the C-terminal end of the pore loop immediately precedes the long flexible linker that runs perpendicular to the membrane plane and connects the pore loop unit to the S6 helix below **(Fig. 3A)**. The S6-proximal interface includes three positions, E942, E945, and D944, where a strong hydrogen bond acceptor or a negative charge is critical for channel function or expression as indicated by the deep mutational scanning scores **(Fig. 3D, Extended Data Fig. 4B)**. In the desensitized state, the side chain of E942 interacts with residues in the S4 helix and S5 helix linker and D944 establishes main-chain interactions with positions 946-948 that would constrain the flexibility of the pore loop, while E945 does not engage in side chain interactions **(Fig. 3A, F)**. We therefore conclude that interactions by residues E942 and D944 are critical for stabilizing the S6-proximal interface in the desensitized state as well as other important states. Although residue E945 does not contribute to stabilizing the desensitized state, it most likely has a stabilizing role in other states, possibly the open state. We speculate that conformational changes during gating may allow E945 to form polar contacts with positively charged residues like R829 and K822 in the S4 helix or even the polar lipid headgroups in the membrane **(Fig. 3F)**. Interestingly, E945 is not conserved in TRPM8 channels from birds, reptiles, or amphibians **(Extended Data Fig. 4C)**, indicating that their energetics of gating at this site might be different from that of most mammalian TRPM8 channels. The mostly negative variant scores at position R950 **(Fig. 3D, Extended Data Fig. 4B)**, which like E945 does not form interactions in the desensitized state **(Fig. 3A, F)**, suggest that its polar character is important for channel function and that this residue may also contribute to stabilizing non-desensitized states, albeit to a lesser extent than E945. Finally, we propose that residue P949 is intolerant of substitutions **(Fig. 3D, Extended Data Fig. 4B)** because the proline is critical for the folding of the pore loop unit relative to the long linker connecting it to the S6 helix **(Fig. 3A)**.

Notably, in semi swapped structures of an avian TRPM8^31^, where the S6 helices and pore loop are rotated by over 50° relative to the rest of the protein, both pore helix and S6-proximal interfaces with the pore loop are formed involving the same sets of critical residues we described in the fully swapped desensitized state structure **(Extended Data Fig. 4E)**. This observation supports the conclusion that the pore loop unit fold and its critical proximal and distal interaction interfaces with the pore helix and S4, S5, and S6 helices are maintained during channel gating. Based on these findings, we propose a mechanism whereby the pore loop unit behaves as a rigid body during channel gating, its position relative to the rest of the complex stabilized by state-dependent interactions within its pore helix and S6-proximal interfaces. In the open state, correct positioning of the pore loop unit is necessary for the ion selectivity filter on its N-terminal end to adopt a conformation that allows ion conduction through the pore.

Finally, in addition to the mechanism we described, it is likely that many mutations in our screen resulted in negative variant scores because they interfered with the folding of the pore loop unit. In particular, residues T930, P938, and L939 are adjacent to C929 and C940, and thus mutations at these sites may interfere with disulfide bond formation during channel folding. Furthermore, we observed that mutations to cysteine at most positions in the pore loop, including at sites with the high or intermediate tolerance of substitutions, resulted in scores that were more negative than the median **(Fig. 3G)**. This suggests that substitutions to cysteine at other sites in the pore loop may result in formation of a disulfide bond involving a distinct cysteine pair than in WT channels, leading to non-functional channels. These observations underscore the importance of the pore loop structure in TRPM8 channel function.

### Structural determinants of gating in the pore of TRPM8

To learn the sequence and structural determinants of gating by the S6 helices in TRPM8, we began by identifying the state-dependence of all side-chain contacts between S6 helix residues 951-974 and the rest of the protein **(Fig. 4A)** in structures reported to represent closed (C1), open (O), and desensitized (D) states^15,30^ **(Fig. 4B)**. We classified each position in this region of the S6 helix into two categories using the cumulative histograms of all variant scores per position: (1) Critical positions that are intolerant of substitutions **(Fig. 4C, D; dark blue, magenta, purple)** and (2) positions with intermediate tolerance **(Fig. 4C, D; grey, lavender, orange, and yellow)**. Based on this classification, the individual variant scores **(Fig. 4E)**, and the structural information **(Fig. 4A, B)**, we divided the S6 helices into five segments for further analysis: the N-terminal end of S6 that connects to the pore loop (gray); the all α-helical region that is in contact with the pore helices and ion selectivity filter (purple, lavender, magenta); the region where a π-helix bulge forms in the open state (orange, yellow); the region that is in contact with the aqueous pore, S5-, S6-, or S4-S5 linker-helices (dark blue).

**Figure 4.**
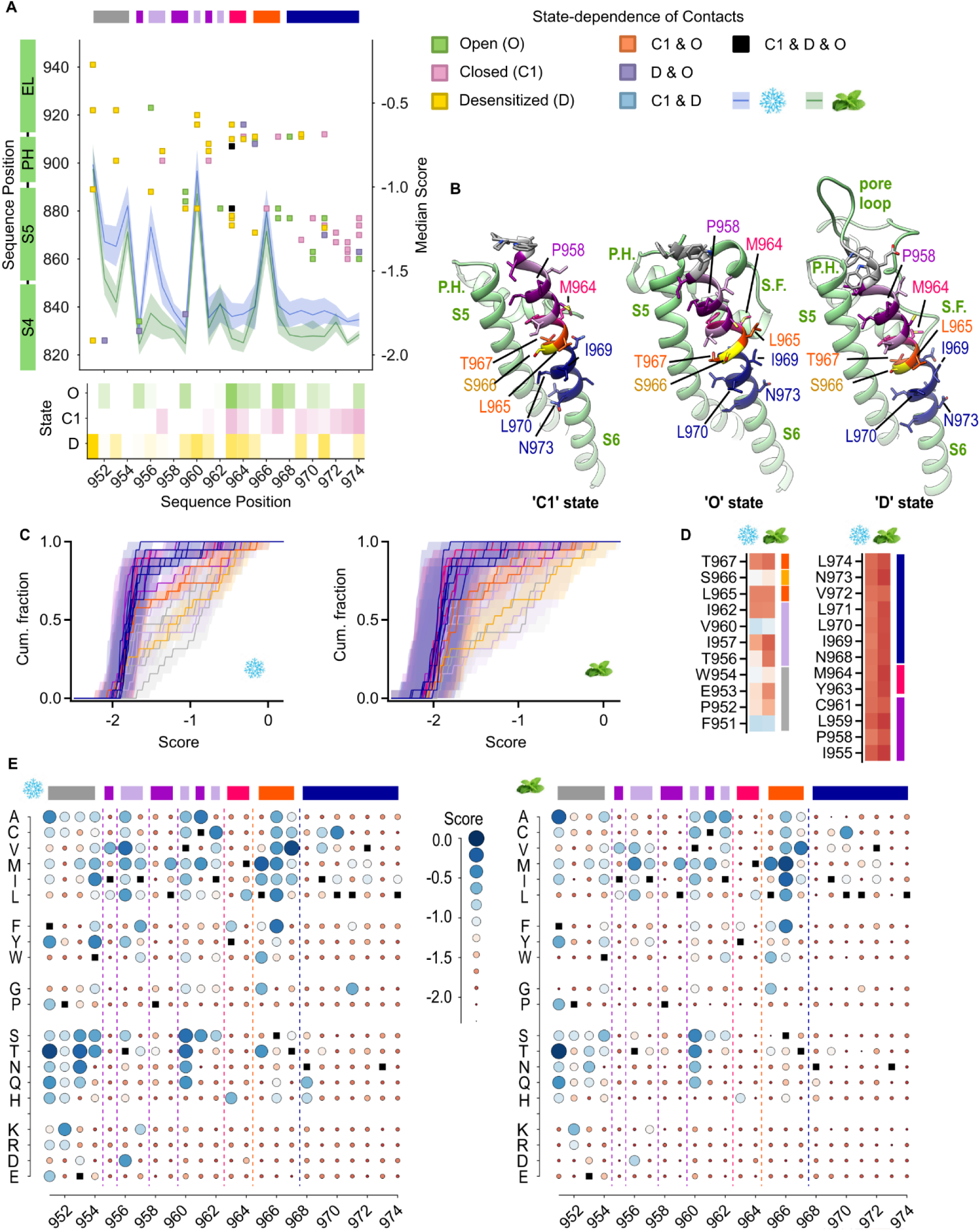
Sequence determinants of gating in the S6 helices of TRPM8. **(A)** Map of residue-residue contacts between S6 helix residues 951-974 (x-axis) and the rest of the protein (y-axis, left) in C1^30^ (PDB: 8E4N), O^30^ (PDB: 8E4L), and D^15^ (PDB: 9B6D) structures of TRPM8. Each color square on the map represents one contact and the color of the square denotes which states have that contact. The heatmap on the bottom insert denotes the number of total atomic contacts by residues 927-974 in each state. The median cold and menthol scores per position are superposed (y-axis, right). The shading is the SEM of the scores across the 19 variants per position. **(B)** Atomic models of the pore of domain of one TRPM8 subunit in C1, O, and D states, with residues 927-974 colored by helix region. **(C)** Cumulative histograms of all variant scores per sequence position between residues 951 and 974, colored by S6 region and tolerance to substitution, as indicated in (D). Shading is the average Lilace^53^ standard error per position. **(D)** Variant classification per S6 helix region and tolerance of substitutions (right-most color bars), showing the median score per position as a color scale. **(E)** Bubble plot of individual variant cold (left) and menthol (right) scores. Black squares denote WT residues, color bars on top denote each of the different S6 helix regions discussed in the text.

The N-terminal portion of S6 formed by residues F951 to W954 is poorly resolved in the C1 and O structures^15,30^ due to the flexibility of this region where the S6 helix becomes partially unwound **(Extended Data Fig. 5A, B)**. Residues F951 to E953 have an intermediate tolerance of substitutions **(Fig. 4C, D)**, and unlike most of the S6 helix, have a preference for small, polar residues without net charge over larger hydrophobic ones **(Fig. 4E, Extended Data Fig. 5C-F)**, which is consistent with the dynamic character of this region and its position at the interface between the solvent and the polar lipid headgroups in the membrane. Residue W954 translocates downward upon channel opening^15,30^ **(Extended Data Fig. 5A-B)**, possibly moving from the polar membrane surface in the closed state into the hydrophobic core in the open state. This gating-dependent change in the polarity of the environment for W954 would explain why this position has greater tolerance for hydrophobic substitutions over hydrophilic ones (**Extended Data Fig. 5C, G**) and suggests that side-chain hydrophobicity at position 954 may contribute to the energetics of channel opening. Finally, based on the strongly negative scores for critical position P958 further down the helix **(Fig. 4C, D, Extended Data Fig. 5C)**, we propose that this proline is required to destabilize the backbone hydrogen bond network at relative position i-4 (i.e. W954) in the helix, favoring unwinding of its N-terminal end during gating as observed in these structures^15,30^ **(Extended Data Fig. 5A, B)**.

The region of S6 between positions I955 and I962 undergoes more limited state-dependent changes, and residues maintain their overall orientation relative to the surrounding S4, S5, and pore helices and ion selectivity filter in C1, O, and D structures^15,30^ **(Fig. 5A)**. We identified two state-dependent interfaces where critical S6 residues in this region form contacts with other helices in either the O (I955, L959; interface with S4 and S5 helices) or C1 and D structures (C961; interface with the pore helix of neighboring subunit) **(Figs. 4A, 5A)**. The intolerance of substitutions at these three positions **(Fig. 5B)** suggests that these residues at these two interfaces have an essential role stabilizing the S6 helix registry while the pore elements that surround this region move significantly during gating. In the C1, O, and D structures residues at positions T956, I957, and V960 face the same cavity behind the pore helix and ion-selectivity filter assembly **(Fig. 5A)**. We propose that their higher tolerance of substitutions **(Fig. 5B)** reflects permissive structural constraints within this interface due to the pronounced structural rearrangements that occur in the surrounding cavity during channel gating. Notably, these three sites have a higher tolerance of conservative substitutions where side chain volume or polarity are similar to the native residues **(Fig. 5B)**. This suggests that stronger perturbations such as the introduction of charges may interfere with channel function by stabilizing or destabilizing certain states through the formation of side-chain interactions between the substituted side chain and polar or charged residues in the pore helices and selectivity filter that are facing this interface **(Fig. 5A)**. In contrast with the three residues above, position I962 was more tolerant of substitutions to hydrophobic than hydrophilic amino acids **(Fig. 5B)**, consistent with its position facing the lipid membrane in the C1, O, and D structures^15,30^ **(Fig. 5A)**.

**Figure 5.**
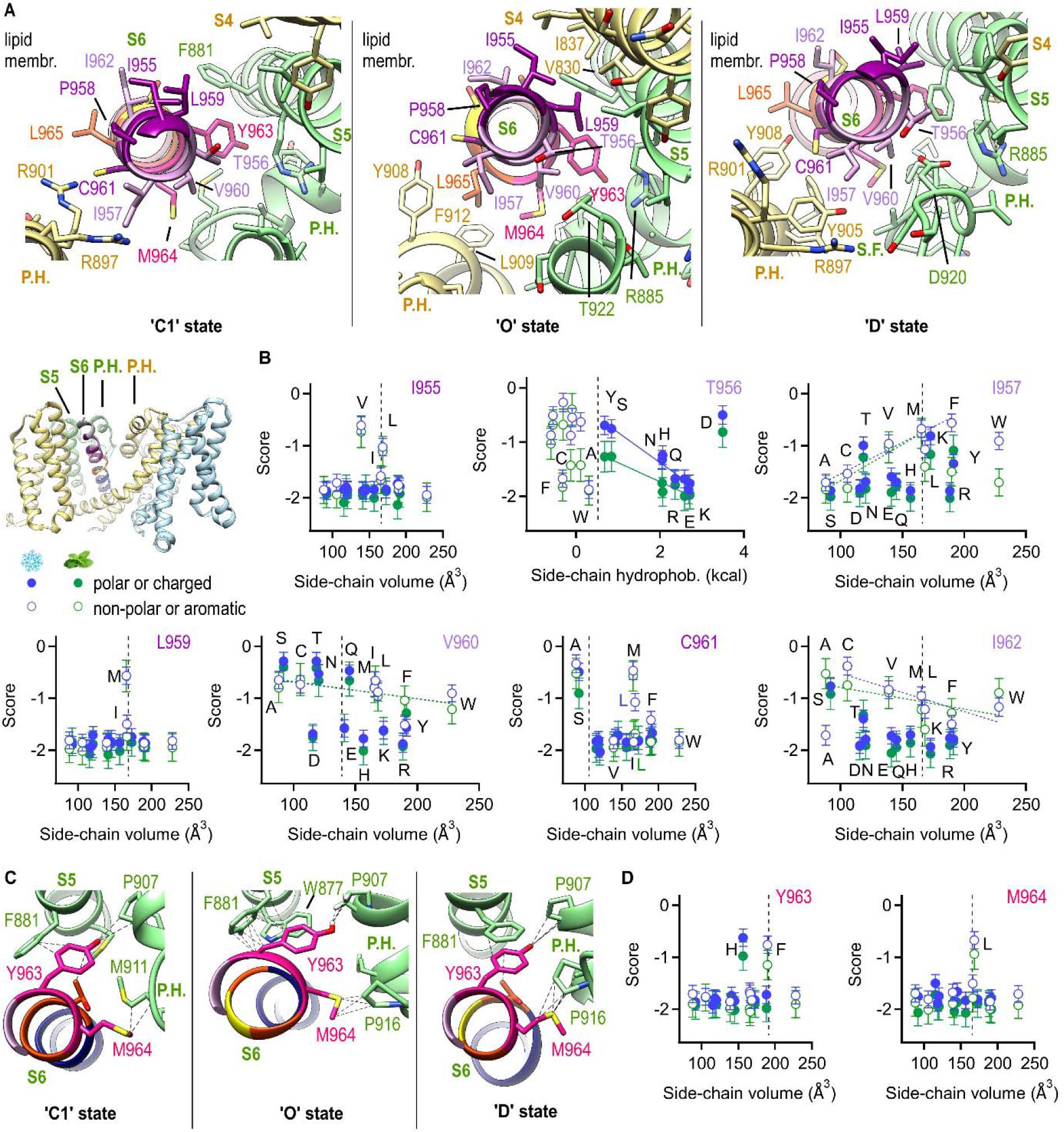
Key determinants within the upper half of the TRPM8 pore. **(A)** Depiction of an S6 helix and adjacent pore elements viewed from above in C1^30^ (PDB: 8E4N), O^30^ (PDB: 8E4L), and D^15^ (PDB: 9B6D) structures of TRPM8. S6 residues I955 to S966 are colored by helix region, the S5, pore helix (P.H.) and selectivity filter (S.F.) of the same subunit colored in green, and the S4 and pore helices of the adjacent subunit in sand. The transmembrane domain in the tetrameric TRPM8 complex in the O state is shown on the bottom left insert, with each subunit shown in a different color. **(B)** Individual variant cold (blue) and menthol (green) scores as a function of substituting side chain volume or hydrophobicity^77^, distinguishing between polar and charged substituting amino acids (closed symbols) or non-polar and aromatics (open symbols). Error bars are the Lilace standard error per variant. Substitutions with proline and glycine were omitted. Dashed vertical lines denote the WT-residue side chain volume or hydrophobicity. **(C)** Structural depiction of the Y963 and M964 interaction interface in C1, O, and D structures, with colors as in (A). Dashed black lines denote non-covalent interactions between the S6 and the rest of the protein. **(D)** Individual variant cold (blue) and menthol (green) scores as a function of substituting side chain volume. Error bars are the Lilace standard error per variant. Substitutions with proline and glycine were omitted. Dashed vertical lines denote the WT-residue side chain volume.

Critical residues Y963 and M964 are localized at the interface between the aqueous cavity and the pore helix and ion-selectivity assembly and establish multiple non-polar interactions with residues in the S5 and pore helices **(Figs. 4A, 5C)**. Notably, within the region of S6 that we examined residue Y963 is the only one that forms interactions with the same two residues (F881, P907) in all three states **(Figs. 4A, 5C)**. Because of these characteristics, and the intolerance of substitutions at positions Y963 and M964 **(Fig. 5D)**, we propose that these residues form non-polar contacts in all states of the channel. These contacts are required to stabilize the upper half of the S6 helix and pore while the helical segments below undergo large changes in registry and secondary structure as a consequence of gating.

Except for the N-terminal end of the helix, the segment between positions L965 and T967 exhibits the largest state-dependent change in helix registry and secondary structure, transitioning from an all α-helical configuration in the C1 and D states to a π-helix bulge in the O state **(Fig. 6A, B)**. The higher tolerance of substitutions at these three positions of the helix relative to the largely intolerant segments immediately above (Y963, M964) and below (N968-L974) is consistent with this region experiencing reduced structural constraints so that it can accommodate the drastic changes in secondary structure associated with gating in this region **(Fig. 4E)**. Furthermore, the individual variant scores at positions L965 and T967 point to a preference for non-polar substitutions of a certain size **(Fig. 6C)**, which suggests that these residues have an important role forming stabilizing interactions in closed and open states as observed in the C1 and O structures **(Figs. 4A, 6B)**. In contrast, residue S966 only shows protein-protein interactions in the C1 and D states but not the O state **(Figs. 4A, 6B)**, where it faces the plasma membrane. Notably, the individual variant scores at position S966 point to a preference for hydrophobic over hydrophilic substitutions **(Fig. 6C)**, suggesting that the hydrophobicity of the side chain at position 966 could contribute to tuning the energetics of channel gating. The rest of the positions after T967 are all intolerant of substitutions **(Fig. 4C-E)**. We propose that each of these positions plays essential roles stabilizing the helix during gating through state-dependent contacts with the S6 helices of other subunits as well as S5 and S4-S5 linker helices **(Figs. 4A, 6A)**.

**Figure 6.**
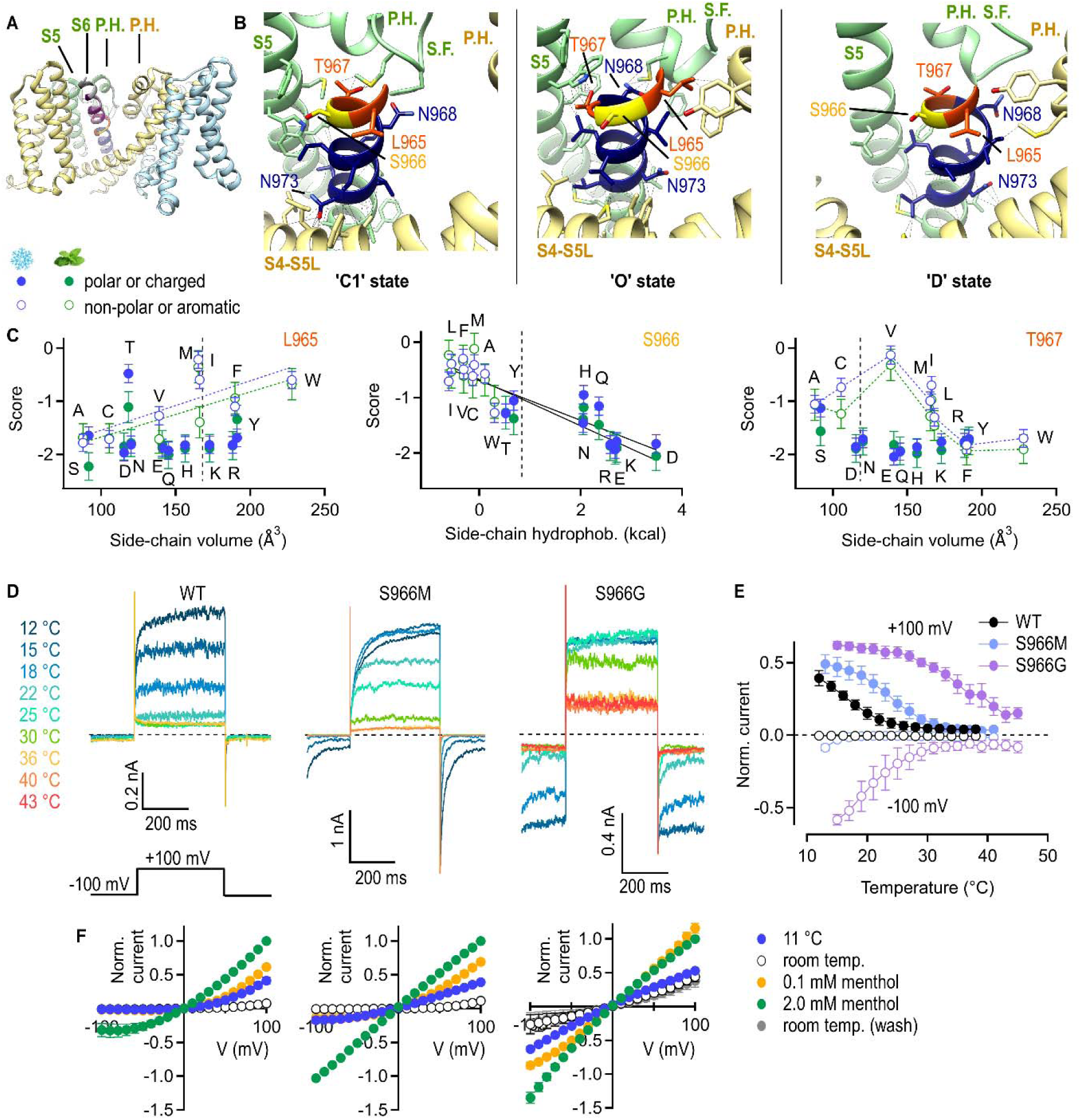
The mechanism of TRPM8 channel gating by the S6 helices. **(A)** Transmembrane domain in the tetrameric TRPM8 complex in the O state with each subunit shown in a different color. **(B)** Depiction of an S6 helix and adjacent pore elements viewed from the membrane side in C1^30^ (PDB: 8E4N), O^30^ (PDB: 8E4L), and D^15^ (PDB: 9B6D) structures of TRPM8. S6 residues L965-N974 are colored by helix region, the S5, pore helix (P.H.) and selectivity filter (S.F.) of the same subunit colored in green, and the S4, S4-S5 linker, S5 and pore helices of the adjacent subunit in sand. Dashed black lines denote non-covalent interactions between the S6 and the rest of the protein. **(C)** Individual variant cold (blue) and menthol (green) scores as a function of substituting side chain volume or hydrophobicity^77^, distinguishing between polar and charged substituting amino acids (closed symbols) or non-polar and aromatics (open symbols). Error bars are the Lilace standard error per variant. Substitutions with proline and glycine were omitted. Vertical dashed lines denote the WT-residue properties. **(D)** Representative current families at ± 100 mV recorded at distinct temperatures denoted by the trace colors from whole cells expressing WT or mutant channels. Dashed lines denote the zero-current level. **(E)** Normalized current-temperature relations obtained from experiments as in (D) for WT and mutant TRPM8. Data shown as mean ± SEM (n = 4). **(F)** Normalized current-voltage relations recorded at each of the conditions denoted by symbol color for WT and mutant TRPM8. Data shown as mean ± SEM (n = 4).

By combining the deep mutational scanning scores with structural information, we have provided a description of the structural determinants of gating in the region of S6 that we examined, where critical state-dependent and independent non-polar contacts stabilize the upper and lower segments of the helix, while formation of a π-helix bulge in the middle segment allows the pore to open. To test whether formation of the π-helix bulge is indeed associated with TRPM8 channel opening, we used patch clamp to evaluate the effect of substitutions to glycine and methionine at position S966 that we predicted would increase flexibility and favor channel opening. Consistently, we found that both mutations resulted in increased channel activity at higher temperatures **(Fig. 6D-E)** and at higher voltages in the presence of menthol relative to WT **(Fig. 6F)**, with S966G showing the strongest effect. We therefore conclude that TRPM8 channel opening in response to cold and menthol involves the formation of a π-helix bulge at the midpoint of the helix. Notably, we made the surprising discovery that except for the formation of the π-helix bulge in the open state, most of the structural and sequence determinants that we described above were also satisfied in the closed and open semi-swapped structures of an avian TRPM8 channel^31^ to the extent that they are resolved **(Extended Data Fig. 6)**. We interpret this as a strong indication of how essential these determinants are for the stability of the S6 helix in the context of the rest of the protein.

## Discussion

In this study we implemented a deep mutational scanning strategy to interrogate the available structural information and determine how TRPM8 channels respond to activators. We found that both cold and menthol have similar effects on the region of the channel that we examined and proposed a gating mechanism whereby the extracellular pore loop functions as a rigid coupling element between the S6 helices and the ion selectivity filter loop. As the S6-, S5-, S4- and pore-helices move during gating, the position of the extracellular pore loop relative to the rest of the pore elements becomes stabilized by state-dependent interactions at the pore helix and S6-proximal interfaces **(see (1), (2), Fig. 7A, C; (5), Fig. 7E)**. The correct positioning of the pore loop, stabilized by these interactions, is essential in the open state of the channel for the ion selectivity filter to conduct ions. In turn, the S6 helices are stabilized during gating by essential interactions that are almost exclusively non-polar and state-dependent, involving distinct interfaces above **(Fig. 7, (6), (7))** and below **(Fig. 7, (9))** the region where formation of a π-helix bulge allows the pore to open **(Fig. 7, (8))**. By benchmarking the scores derived from deep mutational scanning using the Ca^2+^-fluorescence responses of individual variants to cold and menthol, we showed that the variant score distributions at each position in the primary sequence were sufficient to identify critical determinants of channel function or expression. We have utilized this information to delineate the mechanism that we propose, which is supported by strong agreement with the structural data and with the amino acid sequence conservation between species.

**Figure 7.**
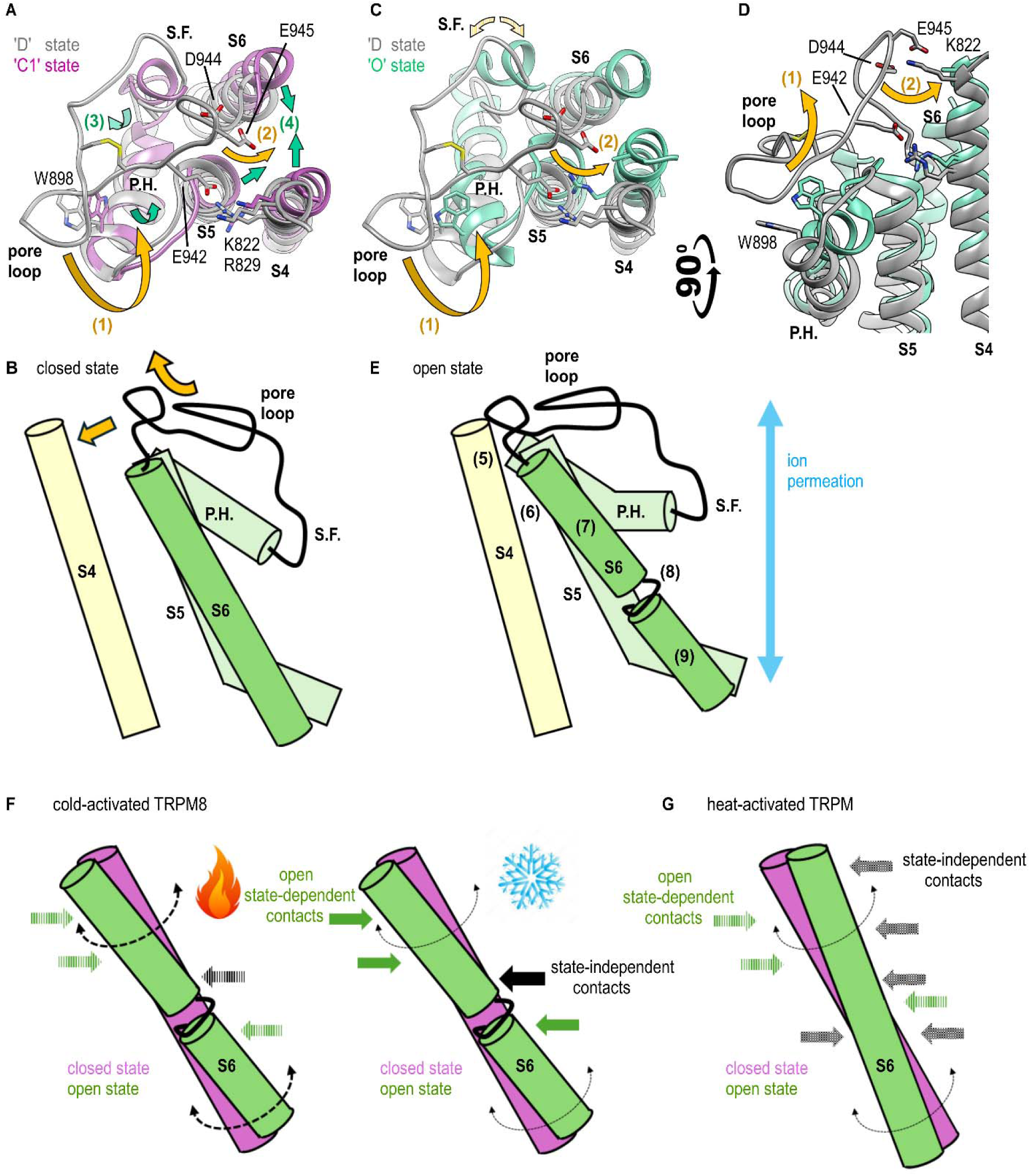
Pore gating mechanism in TRPM8 channels. **(A)** Structural alignment of the pore region in C1^30^ (magenta, PDB: 8E4N) and D^15^ (gray, PDB: 9B6D) structures viewed from above. Arrows denote inferred movements upon channel opening within (1) the S6-distal interface and (2) S6-proximal interfaces of the pore loop, as well as within (3) the pore helices and (4) S4, S5, and S6 helices. **(B)** Cartoon of the TRPM8 pore of one subunit (green) and the S4 helices of the adjacent subunit (sand) in the closed state. Arrows denote the direction of movement of the pore helix during gating. **(C, D)** Structural alignment of the pore region in O^30^ (aqua, PDB: 8E4L) and D^15^ (gray, PDB: 9B6D) structures viewed from above (C) or from the side (D). Dashed arrows denote dynamics within the ion-selectivity filter that are constrained by the pore loop helix, solid arrows denote the same movements as in (A). **(E)** Cartoon of the TRPM8 pore of one subunit (green) and the S4 helices of the adjacent subunit (sand) in the open state. Numbers denote each of the five regions of S6 that are described in the text. **(F)** Cartoon illustrating how open state-dependent (green arrows) and -independent (black arrows) contacts that stabilize S6 helices in the open state of TRPM8 go from weaker (left, dashed arrows) to stronger (right, filled arrows) as the temperature decreases. **(G)** The S6 helices in heat-activated TRPM channels are stabilized by a larger proportion of state-independent contacts, making the helices and pore less dynamic.

Computational methods to predict the effects of mutations on protein function have gained accuracy over the last decade^56^. AlphaMissense scores individual human protein variants based on their predicted pathogenicity on a scale of 0 (benign) to 1 (pathogenic)^57^. Scaling both predicted and experimental scores to be quantitatively comparable showed a qualitatively similar biphasic distribution of scores, except that the distribution of AlphaMissense variants with the most negative scores was much sharper than that of the experimental scores **(Extended Data Fig. 7A)**. Despite these evident differences, we found good overall agreement between predicted and experimental individual variant scores **(Extended Data Fig. 7B-I)**. However, we did observe deviations at multiple positions, including some of the positions that we identified as critical or important (i.e. 930, 935, 939, 942, 945, 950-954) **(Extended Data Fig. 7C-F, asterisks)**. In general, the predicted scores overestimated the tolerance of substitutions at positions within the extracellular pore loop and underestimated that of positions along the entire S6 helix. These observations underscore the importance of obtaining experimental datasets like ours, which can help improve the accuracy of the computational variant effect predictions^58^.

Although initially we used three ‘fully swapped’ structures as a reference to interpret our results^15,30^, we found that the general sequence and structural determinants that we identified within the pore loop and S6 helices were also compatible with a different fully swapped closed state structure of the human TRPM8^31^ **(Extended Data Fig. 6D; ‘C’ state fully swapped)**. We also examined a fully swapped open state structure of the human TRPM8^31^ but were unable to assess its significance due to limited resolution in the pore region and a drastically reduced number of side chain contacts between the S6 helix and the rest of the protein in this structure **(Extended Data Fig. 6D; ‘O’ state fully swapped)**. Notably, we found similar structural constraints involving state-dependent interactions and higher packing density associated with S6 helix positions that were most intolerant of substitutions in either fully swapped or semi swapped closed and open state structures **(Extended Data Fig. 6)**. Importantly, the side chain of S966 interacts with other parts of the channel in closed and desensitized states but faces the hydrophobic lipid tails in the open state in either the fully swapped^59^ or semi swapped^31^ structures. This structural feature agrees with the positive correlation we detected between the hydrophobicity of the substituted side chains and the tolerance of the channel to the substitutions **(Fig. 6C)** and underscores the importance for channel gating of position S966 regardless of whether it involves fully swapped or semi swapped states. However, formation of a π-helix bulge between positions L965 and T967 is unique to the fully swapped open state. We consider that this drastic change in secondary structure in the fully swapped gating model agrees particularly well with our observation that positions 965-967 show a higher tolerance of substitutions than the adjacent portions of the helix **(Fig. 4E)**. It is unclear whether TRPM8 channels can transition between fully swapped and semi swapped conformations when expressed in living cells because of the high energy barrier associated with such a process. However, we conclude that such states of TRPM8 can be observed, at least in cryo-EM grids, because they happen to satisfy most of the structural constraints that we found here to be essential for conferring stability to the S6 helices. Interestingly, unexpected transmembrane helix rearrangements could be seen within the pore domains of prokaryotic voltage-gated channels^60^ whose fold is similar to that of TRP channels. The intrinsic stability of this widely conserved fold is likely what has allowed for the extensive variability in its configurations, dynamics, and associated mechanisms of gating across distinct ion channel families^61,62^.

We observed that most residue-residue contacts formed between the upper half of the S6 helix and the rest of the protein are state-dependent and involve weaker non-polar interactions. We speculate that these weaker hydrophobic interactions can be easily broken, which is required to accommodate the dynamic changes in structure that occur within the S6 helix during gating. Indeed, pore opening in TRPM8 is likely associated with increased structural dynamics in the pore, because this region is poorly resolved in most open state structures^14,30,31^. In addition, TRPM8 is the only TRP channel where a semi swapped conformation of the pore has been observed^31^, further supporting the view that the pore of this channel is particularly dynamic. We propose that the structural dynamics within the TRPM8 pore and S6 helices that occur during gating are also involved in its activation by cold: at higher temperatures the open state of the pore is unstable because the dynamics in the pore overcome the weak and mainly non-polar interactions that can stabilize the S6 helices in their open state **(Fig. 7F, dashed arrows)**. The open state becomes stable at lower temperatures because as the dynamics become attenuated the non-polar interactions formed between the S6 helix and the rest of the protein become sufficiently strong to stabilize the channel in its open state **(Fig. 7F, solid arrows)**. Consistent with this mechanism, measurements of channel activation and deactivation kinetics suggested that cold activates TRPM8 channels by stabilizing an open state, whereas heat activates other TRP channels by destabilizing closed states^7^. Furthermore, based on structural information showing cold-induced conformational changes in the S6 helices^33^ it was proposed that cold activates TRPM8 channels by stabilizing an open state. Evidence from species-dependent differences in TRPM8 cold sensitivity also suggests that in mammals, the pore domain of TRPM8 has an influence on temperature sensing^63–65^. It was also recently shown that regions within the pore of TRPM8 undergo temperature-dependent changes in accessibility to deuterium, further suggesting that the pore may contribute to temperature sensing^31^.

We reasoned that if the dynamic character of the S6 helices in TRPM8 contributes to its ability to open in response to cold, then other TRP channels that share homology with TRPM8 but are instead activated by heat would be predicted to have less dynamic S6 helices constrained by more stable interactions. Consistent with our hypothesis, we found that most of the contacts formed between the S6 helices and the rest of the protein in closed and open structures of the heat-activated^66,67^ TRPM2^68^ and TRPM4^69^ channels were state-independent **(Extended Data Fig. 8A)**. This is in stark contrast with our finding that most residue-residue contacts by the S6 helices of TRPM8 are state-dependent **(Fig. 4A)**, which means that a more extensive reconfiguration of stabilizing interactions is necessary for gating in the cold-activated TRPM8 channel than in its heat-activated homologues **(Fig. 7G)**. Furthermore, we found that the median of the AlphaMissense^57^ pathogenicity predictions per sequence position for the heat-activated^66,67^ TRPM2, TRPM4, and TRPM5 channels showed clear differences relative to the experimental and predicted values for TRPM8 **(Extended Data Fig. 8B)**, suggesting that the structural determinants of gating associated with the S6 helices are different between the heat- and cold-activated proteins. An important aspect of the cold-sensing mechanism that we propose is that although it involves the dynamics of the S6 helices, the interactions of these helices with the rest of the pore elements are as important as the S6 helix itself, such that stimuli acting on other parts of the protein can tune cold sensitivity by influencing the conformation of any of the structural elements that are part of the pore. Finally, it is important to mention that the cold-sensing mechanism we propose does not rule out contributions from other parts of the protein^64,70,71^.

## Methods

### Cloning of expression plasmids

We used the backbone of the promoter-less plasmid KAM05_attB_mCherry, which was a gift from Doug Fowler (University of Washington), to onboard a synthetic gene sequence coding for the rat TRPM8 and an upstream partial Internal Ribosome Entry Site (IRES2) sequence using Twist Bioscience gene synthesis service. We strategically modified this sequence to include or remove restriction enzyme recognition sites without altering the amino acid sequence of the channel, except for the inclusion of a 4 amino acid (Met-Ala-Thr-Thr) insertion at its N-terminus – for clarity we have disregarded this insertion in the primary sequence numbering used in the manuscript. We cloned a synthetic double stranded DNA fragment (Twist Bioscience) coding for GCaMP6s and a downstream partial IRES2 sequence into the plasmid using EcoRI and SacII (New England Biolabs) by restriction/ligation with T4 ligase (New England Biolabs). We included an mCherry fluorophore expressed in frame with TRPM8, which we later used to select cells with an intact reading frame coding for all three proteins. To include the T2A-mCherry sequence downstream of TRPM8 and remove the stop codon of the channel, we used a two-step PCR approach with two pairs of overlap (O_fwd, ctaacatgcggtgacgtcgaggagaatcctggcccaatggtgagcaagggc; O_rev, gtcaccgcatgttagcagacttcctctgccctctttgattttattagcaat) and flanking primers (F1_fwd, ggtgggctgtggcct; F1_rev, aagagctagcctctacttgtagagctcgtccat) and using restriction enzymes MfeI and NheI (New England Biolabs) for restriction/ligation cloning. The pCAG-NLS-HA-Bxb1 plasmid was purchased from Addgene (plasmid #51271).

To generate the acceptor plasmid for the library, we used restriction sites MfeI and SphI, which flank the sequence coding for the transmembrane domain of TRPM8, to clone a double stranded DNA insert (Twist Bioscience) that included BsmBI sites flanking the variable-sequence region of the library from positions 927 to 974. Plasmids coding for mutants Y745A, H846A, R842A, and K856A and each of the four synonymous mutations (L843, CTC to TTG; L871, TTG to CTC; S902S, TCT to AGC; V983, GTG to GTC) we constructed by restriction/ligation cloning of synthetic DNA fragments (Twist Bioscience) using MfeI and SphI, while BsmBI sites in the acceptor plasmid were used for cloning synthetic DNA fragments coding for each of the 28 single mutants used for benchmarking the scores. Cloning design and plasmid map curation were done with SnapGene, and all constructs were verified using Sanger (EtonBio) and/or full plasmid sequencing (PlasmidSaurus).

### Plasmid library generation

A library of all possible 5,472 single-residue variants and 288 single residue deletions within the transmembrane (TM) domain (positions F735-Y1022) of the rat TRPM8 was designed using the SPINE script^51^. Briefly, this sequence spanning 288 amino acids was divided into six segments of 48 amino acids each, and SPINE was used to define all variant DNA coding sequences within each of the segments. A single codon per amino acid was included for each position in the library. Appropriate BsmBI enzyme recognition sites flanking both ends of the variant coding sequences were included by SPINE to maintain the open reading frame after cloning of the segments into the rest of the expression plasmid by the Golden Gate method (see below). SPINE also included segment-specific primer binding sites on each end of the library sequences to selectively amplify only variant sequences within each of the six segments. These sequences were used to obtain a pooled 230 nucleotide oligo library of < 7,500 variants synthesized by the Agilent SurePrint Oligo Synthesis service. Notably, Agilent provided a Hi-Fidelity library free of charge, which is what was used in this project.

From the pooled oligo library, a subset of sequences coding for variants between positions 927-974 was amplified using Platinum SuperFi II 2x PCR Master Mix (Invitrogen), 200 nM of each forward and reverse primers, and 15 amplification cycles with an annealing temperature of 57. The specificity of the reaction was verified by agarose gel electrophoresis, and the amplified oligos were purified using the Oligo Clean & Concentrator kit (Zymo Research) and an elution volume of 15 μL of 60 °C nuclease-free water, and their concentration after purification was measured on a fluorescent plate reader using PicoGreen reagent (Invitrogen) and 96-well plates. The amplified DNA fragments were then cloned into the TRPM8 acceptor plasmid that included appropriate BsmBI sites flanking the segment of variable sequence. The Golden-Gate reaction included 0.6 µL T4 ligase (NEB), 2 µL T4 ligase buffer (NEB), 1.2 µL BsmBI (NEB), 80 ng DNA fragments, and 500 ng TRPM8 acceptor plasmid. The reaction temperature was cycled 60 times between 42 for 5 minutes and 16 for 10 minutes, after which temperature was held at 42 for 20 minutes followed by heat-inactivation at 80 for 10 minutes. The reaction product was purified using the Zymo Clean and Concentrator-5 kit (Zymo Research) as per the manufacturer’s protocol and eluted in 10 uL of 60 nuclease-free water. A volume of 2 uL of eluant was electroporated into E. cloni 10G Elite electrocompetent cells (Biosearch Technologies) following manufacturer’s instructions. After electroporation, a volume of electroporated cells containing between 1-2 x 10^5^ unique transformants was inoculated into 0.1 L of Lysogeny Broth (LB) liquid media supplemented with 100 μg/mL ampicillin and incubated at 37 °C for 18 hours with shaking. Dilutions of the media at the time of inoculation were plated on ampicillin agar dishes to confirm the number of transformants. Plasmids were extracted 18 hours after inoculation using the Qiagen Miniprep Kit as per the manufacturer’s instructions, using nuclease-free water at 60 for elution.

The plasmid library was assessed by next-generation sequencing on a NovaSeq platform (Illumina). Plasmids were prepared for NGS by digestion with MfeI and SphI (NEB) for 5 hours. The digestion reaction was then run on a 1% TAE gel and the band corresponding to the transmembrane domain sequence region was cut out and purified using the QIAquick Gel Extraction Kit (Qiagen) as per the manufacturer’s instructions and eluted in 60 nuclease-free water. The fragments containing the Transmembrane Domain region were sent to the UT Austin Genomic Sequencing and Analysis Facility for sequencing. Variants were called using the satmut-utils python pipeline ^52^, which showed homogenous coverage of the missense variants in the library **(Extended Data Fig. 1C)**.

### Tissue culture and protein expression in landing pad cells

Proteins were expressed using HEK293-derived iCasp9 landing pad cells^46^, which were a generous gift from Kenneth Matreyek (Case Western Reserve University). Cells were initially kept under adhesion conditions at 37°C with a 5% CO_2_ atmosphere in high-glucose DMEM media supplemented with pyruvate, L-glutamine, phenol red, 10 % (v/v) heat-inactivated fetal bovine serum, 10 μg/mL gentamicin, and 2 μg/mL doxycycline. Working stock solutions of AP1903 in DMSO were prepared at 10 μM from an initial stock solution of 10 mM. The doxycycline working stock was prepared at a concentration of 2 mg/mL using sterile water. Recombination of the expression plasmids for WT or individual TRPM8 variants into the landing pad was carried out as follows: Day 1, iCasp9 cells were seeded onto 6-well plates pre-treated with 6 μg/mL bovine plasma fibronectin (Sigma) prepared in PBS, using a single well per construct, transfected with the Bxb1 plasmid using Fugene HD (Promega) following the manufacturer’s instructions, and kept in complete media without doxycycline. Day 2, cells were transfected with the promoter-less plasmids. Day 3, media was supplemented with doxycycline to induce expression of landing pad genes. Day 4, media was supplemented with doxycycline and 10 nM AP1903 (MedChemExpress) for negative selection of unrecombined cells. Unrecombined cells die because they express the inducible Caspase 9 (iCasp9) that becomes active upon AP1903-induced oligomerization, whereas recombined cells survive because they no longer express iCasp9. Day 5, cells were washed multiple times with media supplemented with doxycycline and left to recover.

Two to three days after negative selection, cells in each well were transferred to tissue-culture-treated T25 flasks, and upon reaching confluency were transferred into suspension growth conditions; cells were detached using 0.25% trypsin and resuspended in FreeStyle 293 Expression Medium (Gibco), supplemented with 2% (v/v) heat-inactivated fetal bovine serum, 10 μg/mL gentamicin, and 2 μg/mL doxycycline. After resuspension, the cells were washed once with suspension growth media and then transferred to 125 mL baffled-bottom glass flasks. Suspension cultures were maintained at 37 C with an 8% CO_2_ atmosphere and constant shaking at 135 rpm using a Celltron shaker (Infors HT). Cell density was maintained between 0.5 and 2×10^6^ cells/mL, and cultures were resuspended in fresh media and transferred to a clean flask every other day to minimize cell aggregation. A Countess II FL (Thermo Fisher) with GFP and mCherry filter cubes was used to count cells during culture.

Mutants Y745A, H846A, R842A, K856A, and the four synonymous variants were recombined and analyzed once. For the 28 individual mutants used to benchmark the scores, three independent recombination reactions were carried out, and cell lines from each recombination were cultured and analyzed separately to obtain three fully independent biological replicate experiments per mutant. Each recombination involving single variants always included a well of cells recombined with WT TRPM8 plasmid as a control.

To express the plasmid library, all wells in a fibronectin-treated 6 well plate were seeded with iCasp9^46^ cells at 50% confluency, and recombined as described above. When recombining the libraries a plasmid transfection mix composed of 3 μg of library plasmid DNA, 300 μL Opti-MEM I 1X (Gibco), and 6 μL of FuGENE HD transfection reagent (Promega) was used. After recombination and cell recovery from the negative selection, each well was transferred into a separate T25 tissue-culture treated flask, and upon reaching confluency the contents of each flask were transferred into suspension growth conditions as described above. Library cultures were kept in one to four 125 mL baffled bottom glass flasks, each with up to 40 mL of suspension media. Two separate library recombination reactions were carried out, and each resulting cell library culture was sorted separately to obtain the two independent biological replicates reported here.

### Flow cytometry and cell sorting

To analyze the Ca^2+^-fluorescence response to room temperature, cold, or menthol (0.2 and 2 mM) of cells expressing WT TRPM8 or individual channel variants, aliquots of 10 x 10^6^ cells per variant that were kept in suspension were pelleted at 1,300 x g, washed once with an extracellular solution composed of 130 mM NaCl, 4 mM KCl, 10 mM HEPES, 10 mM glucose, 40 mM sucrose, and 0.2 mM CaCl_2_ (pH 7.4), and resuspended in 1 mL of the same solution. For each variant, the cell suspension was distributed into four 5 mL polystyrene round bottom tubes. An additional 0.25 mL extracellular solution was added to the room temperature and cold samples, whereas 0.25 mL 2X menthol solution (0.2 or 4 mM) were added to the menthol samples right before analysis. Cold samples were kept on ice for 5-10 minutes before analysis. Samples were analyzed on a BD FACS Aria Fusion SORP Cell Sorter (BD Biosciences) at the Center for Biomedical Research Support Microscopy and Flow Cytometry facility at UT Austin. Single live cells were gated based on their forward and side-scatter (FSC-A x SSC-A and FSC-A x FSC-H), and the GCaMP6s fluorescence of 10 x 10^3^ mCherry^+^ cells were recorded. The resulting fcs files were opened using Floreada software (http://floreada.io) and saved as .csv files for further analysis using Igor Pro 9.

The analysis in triplicate of the 28 single mutants used to benchmark the scores was carried out in distinct batches starting from the recombination process, each including a sample of cells expressing WT TRPM8. GCaMP6s fluorescence histograms were obtained for each sample, and the x-axis scaling of the WT TRPM8 histograms in each batch was adjusted such that their peaks were aligned at each of the four experimental conditions. The x-axis scaling of the fluorescence histograms of the mutants was adjusted by the same amount as the WT cells from their corresponding experimental batch. The resulting mean ± SEM histograms are displayed on **Extended Data Fig. 3C**.

For sorting and analysis of the cell library at least 160 million cells were kept in suspension on the day of sorting. On that same day, library suspension cultures were supplemented with 1% of cells expressing each of the four synonymous variants. Before sorting, 10 x 10^3^ cells expressing the synonymous variants, as well as aliquots of the library, were prepared as described above and analyzed at room temperature or after cold stimulation to ensure that the distribution of GCaMP6s fluorescence was consistent with previous measurements. Cell-sorter gain and cell-sorting gates were adjusted based on these observations to include comparable numbers of cells between replicates in each of the groups of low, medium, and high GCaMP6s fluorescence. During initial development of the assay, we determined that the GCaMP6s fluorescence intensity remained elevated after stimulation with cold or menthol in cells expressing WT TRPM8 for at least 30 minutes. Therefore, to completely rule out any time-dependent biases in cell fluorescence during sorting, each library sample tube was prepared as described for the individual variants, and cells were sorted for no longer than 15 minutes per sample tube before being exchanged with a freshly prepared sample. Single live cells were gated based on their forward and side-scatter (FSC-A x SSC-A and FSC-A x FSC-H), and mCherry^+^ cells were sorted into groups of low, medium, and high GCaMP6s fluorescence. The total number of sorted cells per group, experimental condition, and replicate number are included in Experimental Data Table 1. Importantly, cell sorting in each of the low, medium or high GCaMP6s groups was stopped upon reaching 5 x 10^5^ cells. Cells were collected into 5 mL polypropylene round-bottom tubes.

### DNA amplicon library generation

After sorting, all collection tubes containing cells from the same low, medium, or high GCaMP6s cell-sorting groups were combined into a single 15 ml polypropylene conical tube. Each 15 ml polypropylene conical tube was centrifuged at 1,300 x g for 5 minutes. The supernatant was then aspirated from the pellet, and samples were frozen at -20°C until further processing. Genomic DNA (gDNA) extraction and purification was carried out for each of the cell sample pellets using the DNeasy Blood and Tissue kit (Qiagen) following manufacturer’s instructions and eluted into 0.2 mL of nuclease-free water. Aliquots of each purified sample were submitted to the Genomic Sequencing and Analysis Facility (GSAF) for quantitation of the DNA concentration by Qubit Fluorometer. Yields from this step depended on the amount of cells collected for each GCaMP6s-intensity group and ranged from 3.8 to 70.2 ng/μL.

DNA amplicons were generated via PCR using a reverse primer that binds within the genomic landing pad downstream from the recombined plasmid sequence (Fowler_REV_Illumina, GTCTCGTGGGCTCGGAGATGTGTATAAGAGACAGCTCTTCGCCCTTAGACACCA) and a forward primer that binds upstream of the sequence coding for the transmembrane domain region of TRPM8 (TM_BT_GEN_FWD, GAATATCCTGTGTCTGTTCATCATC). The entirety of the genomic DNA eluate per sample was distributed across 24 identical reactions. Each reaction thus contained 25 uL Platinum SuperFi II 2x Master Mix (Invitrogen), 2 μL MgCl_2_, 1 μL of each primer 10 μM working stock, 30-600 ng of gDNA template, and enough nuclease-free water to make a total volume of 50 μL. PCR reactions were denatured at 95 for 15 seconds, annealed for 15 seconds at 61, and elongated at 72 for 2 minutes for a total of 29 cycles. All PCR reactions from a single sample were combined into a 15 ml conical tube and purified using the Zymo Clean and Concentrator-5 kit. Briefly, binding buffer was added in a 5:1 volume ratio to the pooled PCR sample and mixed by vortexing for 2 seconds. The mixture was distributed into 3 spin columns with collection tubes, and the DNA purified following manufacturer’s instructions using 25 μL of 60 DNase/RNase-free water for elution. The elution step was repeated with an additional 10 uL of 60 water. The eluate was digested with MfeI and SphI (NEB) for 3 hours at 37 . Following digestion, samples were run on a 0.66% agarose gel made with TAE buffer and the 1 kb band corresponding to the transmembrane domain coding sequence was cut and purified using the QIAquick Gel Extraction Kit (Qiagen) using an initial elution volume of 20 μL followed by a second elution with 10 μL 60°C nuclease-free water. Samples were then sent to the GSAF for further processing and sequencing.

### Next generation sequencing of libraries

Libraries were generated from the DNA amplicons at the UT GSAF core facility. Qubit quantification and an Agilent Bioanalyzer were used to assess the size and concentration of the amplicons prior to library preparation. The roughly 1,200 base pair (bp) amplicons were then mechanically fragmented on a Covaris S2 instrument using 6 cycles of 25 seconds, 10% duty cycle, intensity 5 and 200 cycles per burst, to produce fragments of approximately 300-500 bp. DNA libraries were constructed using the NEBNext Ultra II DNA library prep kit (NEB), using full-length Illumina adapters containing unique dual indices for the i5 and i7. Final libraries were then assessed for quality and concentration with the Agilent HS DNA kit and qPCR with the Kapa SYBR Fast kit (Roche). Sequencing was completed on the NextSeq1000, P2 300 cycle kit, paired end 150. The final loading concentration was 540 pM, producing approximately 25 million reads per sample. **(Extended Data Table 1)**.

### Sequence data analysis

Raw reads were processed using the satmut-utils python pipeline^52^. Nucleotide sequences containing the variant calls were reconstructed using the reference sequence and translated into amino acid sequences using the standard codon table. Amino acid sequences having more than one amino acid change, along with synonymous mutations which were not among the intentionally included list, were filtered out, and a table cataloguing the frequencies of all amino acid missense alterations at each codon was produced from the full set of single-amino-acid variant sequences.

Variant effect scores were generated using Lilace version 1.0.0 ^53^, a Bayesian hierarchical model for fluorescence activated cell sorting-based deep mutational scanning data. Lilace estimates the effect size of unimodal fluorescence distribution shifts for nonsynonymous variants relative to a synonymous variant reference based on the NGS read counts per variant across different groups of cells that were sorted by their fluorescence intensity. Three sets of individual variant scores and corresponding standard errors were obtained per experimental condition – one set of scores obtained from the read counts from each of the two replicates separately, and one combined set where the read counts from each of the two replicates contributed to the score. Except for the correlation plots between replicates, all scores displayed in the manuscript are the combined scores obtained from read counts from both replicate experiments. The number 123 was used as seed for all score calculations.

As synonymous variant references, we used the four silent mutation constructs that we combined with the library cells on the days in which libraries were sorted: L843, CTC to TTG; L871, TTG to CTC; S902S, TCT to AGC; V983, GTG to GTC. The number of cells that we sorted into each of the three groups of low, medium, and high GCaMP6s fluorescence intensity was mostly similar between groups and independent of how many cells in the library were present in each of the three groups **(Extended Data Table 1)**. We also obtained a comparable number of NGS reads from each of the three groups of sorted cells **(Extended Data Table 1)**. In Lilace, scaling factors π_L_, π_M_, and π_H_ incorporate information about the proportion of library cells within each of the three GCaMP6s fluorescence intensity intervals: Replicate 1 cold, π_L_ = 0.01198939, π_M_ = 0.6477886, π_H_ = 0.340222; Replicate 1 menthol, π_L_ = 0.3571429, π_M_ = 0.3970588, π_H_ = 0.2457983; Replicate 2 cold, π_L_ = 0.08559977, π_M_ = 0.5751503, π_H_ = 0.33924991; Replicate 2 menthol, π_L_= 0.18537074, π_M_= 0.5621242, π_H_= 0.25250501. Lilace’s positional prior was disabled so that effect sizes at each position could be estimated without pooling information across individual variants of the same position. Subsequent analyses and visualization of variant effect scores were performed in Igor Pro 9 (WaveMetrics) and Microsoft Excel.

The small number of synonymous variants that we included, and the larger fraction of reads found for each of the synonymous variants relative to the missense variants, caused the score distribution that we obtained to be shifted towards more negative values. Therefore, the inferred score distribution of WT-like variants in the library is centered at a value close to -0.5 instead of 0. To provide an accurate reference to interpret the variant effect scores, we benchmarked the variant scores using the GCaMP6s fluorescence response profiles to cold and menthol that we individually measured for 28 variants.

### Contact map generation

Data processing, analysis, and visualization was performed in version 4.3.3 of R (R Core Team 2024) using the *tidyverse* version 2.0.0 suite of packages. Atomic coordinates for each structure of interest were retrieved from the RCSB Protein Data Bank and opened using *bio3d* version 2.4-4^72^. For contact calculation, we devised a function that emulates UCSF ChimeraX’s^73^ built-in *contacts* function which determines valid contacts between atomic pairs based on a calculated overlap value; briefly, each atom in the structures is assigned a Van der Waals (VDW) radius based on Bondi’s mean VDW radii^74^. This is necessary because PDB files do not contain any molecular geometry data which is required for using United-Atom VDW radii^75^, which ChimeraX automatically determines through the program IDATM^76^. We also calculated pairwise Euclidean distances between all atoms and subtracted these values from their corresponding VDW radii to obtain an overlap value. Just as done with the ChimeraX *contact* function, atomic pairs are only considered to be a valid contact if the overlap is below 0.6 and above -0.4. Valid contacts between side-chain atoms are then pooled per position and subsequently used for visualizing residue-residue interactions. We created contact maps per state by plotting contacts along S6 helix positions 927-974 in one subunit and positions spanning from the S4 to the S6 helix in all subunits. Heatmaps were obtained by summing across all atomic contacts per residue from position 927 to 974, where color intensity represents the number of atomic contacts at each position relative to the total contacts per state.

### AlphaMissense predictions and data scaling

AlphaMissense^57^ pathogenicity predictions for human TRPM8, TRPM2, TRPM4, and TRPM5 were downloaded from the AlphaMissense home website (https://alphamissense.hegelab.org/). Because these scores are defined between 0 and 1, all predictions were scaled by a factor of -2. To perform quantitative comparisons between the scaled AlphaMissense predictions and the experimental data, cold and menthol score distribution histograms were generated, and their x-scaling adjusted such that the peak of the negative score distribution was centered at -2, and the much broader peak corresponding to the rest of the scores was centered at 0. AlphaMissense scores are included in the **Supplementary Data 2** file.

### Patch-clamp electrophysiology

Cells grown in suspension were plated into 35 mm plastic dishes each containing a 20mm glass coverslip and allowed to attach to the glass for 30 minutes before starting experiments. Data was acquired at 5 kHz, filtered at 1 kHz, and digitized using a dPatch amplifier and SutterPatch software (Sutter Instrument). Pipettes were pulled from borosilicate glass (1.5 mm O.D. x 0.86 mm I.D. x 75 mm L; Harvard Apparatus) using a P-97 puller (Sutter Instrument) and heat-polished to final resistances between 0.2 – 1.0 MΩ using a MF-200 microforge (World Precision Instruments). An agar bridge (1M KCl; 4% w/v agar; teflon tubing) was used to connect the ground electrode chamber and the main recording chamber. Recordings were carried out in a temperature-control Delta T imaging chamber (Bioptechs), and solutions containing menthol were delivered using the gravity-fed RSC-200 motorized perfusion system (BioLogic). The Delta T chamber enables uniform and rapid temperature changes by delivering heat directly from the cover glass floor of the chamber, which has an electrically conductive coating on the underside opposite to where cells are located. For cooling, a metal ‘cooling ring’ (Bioptechs) tube shaped into a loop that fits within the Delta T recording chamber was used to rapidly reduce temperature by flowing ice-water through the ring. All experiments were done in the whole-cell configuration of the patch clamp technique, in which cells were lifted from the glass and placed millimeters away from a bead thermistor probe used to record the temperature in the chamber during experiments. The bath solution consisted of 130 mM NaCl, 10 mM HEPES, 10 mM EGTA, and 40 mM sucrose (pH 7.4). This same solution supplemented with 10 mM MgCl_2_ was used in the pipette. A 0.2 M menthol stock solution prepared fresh in DMSO was used to make a 0.1 M solution containing 50% (v/v) DMSO and 50% (v/v) recording solution, which was used as a working stock.

Current-voltage (I-V) relations were obtained by stepping the voltage from -100 to +100 in 10 mV increments for 200 ms per test pulse. Voltage was held at -60 mV for 200 ms before and after each test pulse, and voltage was held at 0 mV for 2 s between pulses. I-V relations were normalized to the mean steady-state current magnitude at 100 mV in the presence of 2 mM menthol. The voltage stimulation protocol described above was also used for measuring the menthol concentration-response relations, but only the curves at ± 100 mV are shown. Current-temperature (I-T) relations were obtained by stepping voltage between +100 and -100 mV for 400 ms per pulse. Voltage was held at 0 mV for 1.8 s and stepped down to -100 mV for 200 ms before each pair of test pulses. These experiments were started at room temperature, where a control I-V relation was acquired. While pulsing between ± 100 mV as described above, temperature was increased to 30-45 °C and the I-T relations were acquired as temperature was lowered to ∼10 °C. An I-V relation was measured at 10 °C. Temperature was increased back to 22 °C (room temperature), where I-V relations in the presence of 0.1 and 2 mM menthol were acquired, followed by a final I-V relation in control solution without menthol. I-T relations were normalized to the mean steady-state current magnitude at 100 mV in the presence of 2 mM menthol. All analysis was carried out using Igor Pro 9 (Wavemetrics) or Microsoft Excel.

### Sequence alignments

Sequence alignments were generated using EMBL-EBI Clustal Omega and displayed using Jalview.

## Acknowledgements

We thank Douglas Fowler (University of Washington) and Kenneth Matreyek (Case Western Reserve University) for their gifts of the KAM05_attB_mCherry plasmid and the iCasp9 landing pad cell line, as well as for sharing the iCasp9 recombination protocol and providing helpful discussions and advice at the start of this project. Cell sorting was carried out at the University of Texas at Austin Center for Biomedical Research Support Microscopy and Flow Cytometry Facility (RRID:SCR_021756). Next-generation sequencing was performed at was performed by the Genomic Sequencing and Analysis Facility at UT Austin, Center for Biomedical Research Support (RRID:SCR_021713). We thank Helen Chen for technical assistance, and Eric Senning (UT Austin) and Willow Coyote-Maestas (UCSF) for helpful discussions. We thank Cailin Mulry, Renee Stephenson, Joshua Chiang, Afroza Khan, and Riya Bansal for commenting on the manuscript. This work was supported by UT Austin startup funds and a NINDS R00 grant to A.J.O. (4R00NS101053).

## Author Contributions

A.T. conceptualized the project, generated essential protocols and reagents, implemented initial bioinformatic analysis routines. A.P. and V.L. generated essential protocols, obtained experimental data, curated and organized data, contributed to data visualization and analysis. C.G. and A.V. contributed to generating essential protocols and reagents. R.S. contributed to all experiments involving flow cytometry. M.B. and D.W. contributed to bioinformatics analysis – variant calling and variant scoring. J.P. contributed with all experimental procedures involving NGS. I.H. and C.C. contributed to generating the initial variant calling procedure. A.J.O. conceptualized and administered the project, acquired funds, performed electrophysiology experiments, and contributed to data analysis and curation, visualization, and drafting the first manuscript. All authors contributed to reviewing and editing the manuscript.

## Competing interests

The authors declare no competing interests.

## Materials & Correspondence

Corresponding author: Andrés Jara-Oseguera

Material availability: Please contact the corresponding author for any materials requests or questions.

## Data Availability

All electrophysiology and fluorescence data are displayed on the manuscript figures. Plasmids used in this study will be made available upon request. The scaled AlphaMissense predicted pathogenicity scores and individual variant scores and standard errors are included for each replicate experiment as well as the combined replicate scores for cold and menthol as **Supplementary Data 1**. Raw read counts for each variant detected within each of the low, medium, or high GCaMP6s fluorescence groups per replicate and experimental condition are included as **Supplementary Data 2**. The DNA oligo sequences of the variant library from positions 927 to 974 of the rat TRPM8 are included as **Supplementary Data 3**. All custom scripts utilized in this study can be found at https://github.com/Jara-Oseguera-Lab/rTRPM8_DMS.

---

**Extended Data Figure 1.**
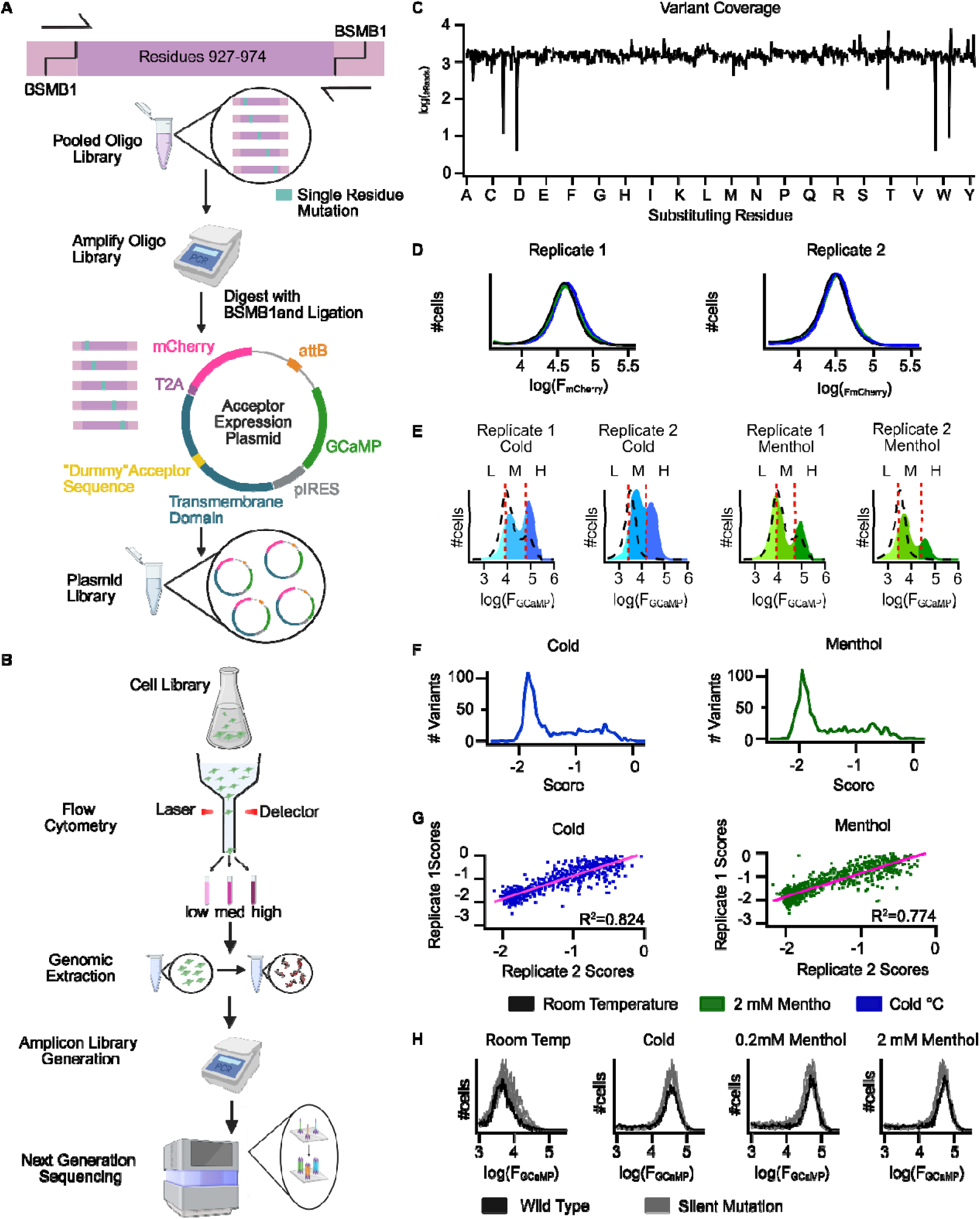
A deep mutational scan of the pore of the TRPM8 channel. **(A)** Cartoon representation of the experimental strategy used in this study. Libraries were generated from a pooled oligo library containing fragments coding for each type of single-residue missense variant within positions 927-974 of the rat TRPM8 channel. Oligo fragments were PCR-amplified and cloned into ‘acceptor’ expression plasmids by Golden Gate cloning using BsmBI sites^51^. **(B)** The plasmid library was expressed in iCasp9 cells^46^, and cells expressing mCherry were sorted by flow cytometry into groups of low, medium, or high GCaMP6s fluorescence intensity after stimulation with cold or 2 mM menthol. The genomic DNA was extracted from the sorted cells, from which the TRPM8 coding region was amplified by PCR, cleaved by a restriction endonuclease to reduce amplicon size, and prepared for next-generation sequencing. **(C)** NGS read counts per variant in the plasmid library, showing homogenous variant representation. **(D)** mCherry fluorescence intensity distribution in the library cells that were sorted. **(E)** GCaMP6s fluorescence intensity distribution of the library cells, measured at room temperature (dashed black curves, not sorted), or after stimulation with cold (blue) or 2 mM menthol (green) during cell sorting. Dashed red lines denote the three cell-sorting bins. **(F)** Deep mutational scanning score distribution for cold and menthol experiments. **(G)** Correlation between individual variant scores from two biological replicates. **(H)** GCaMP6s fluorescence responses in four silent mutations included in the library (gray traces) relative to cells expressing WT.

**Extended Data Figure 2.**
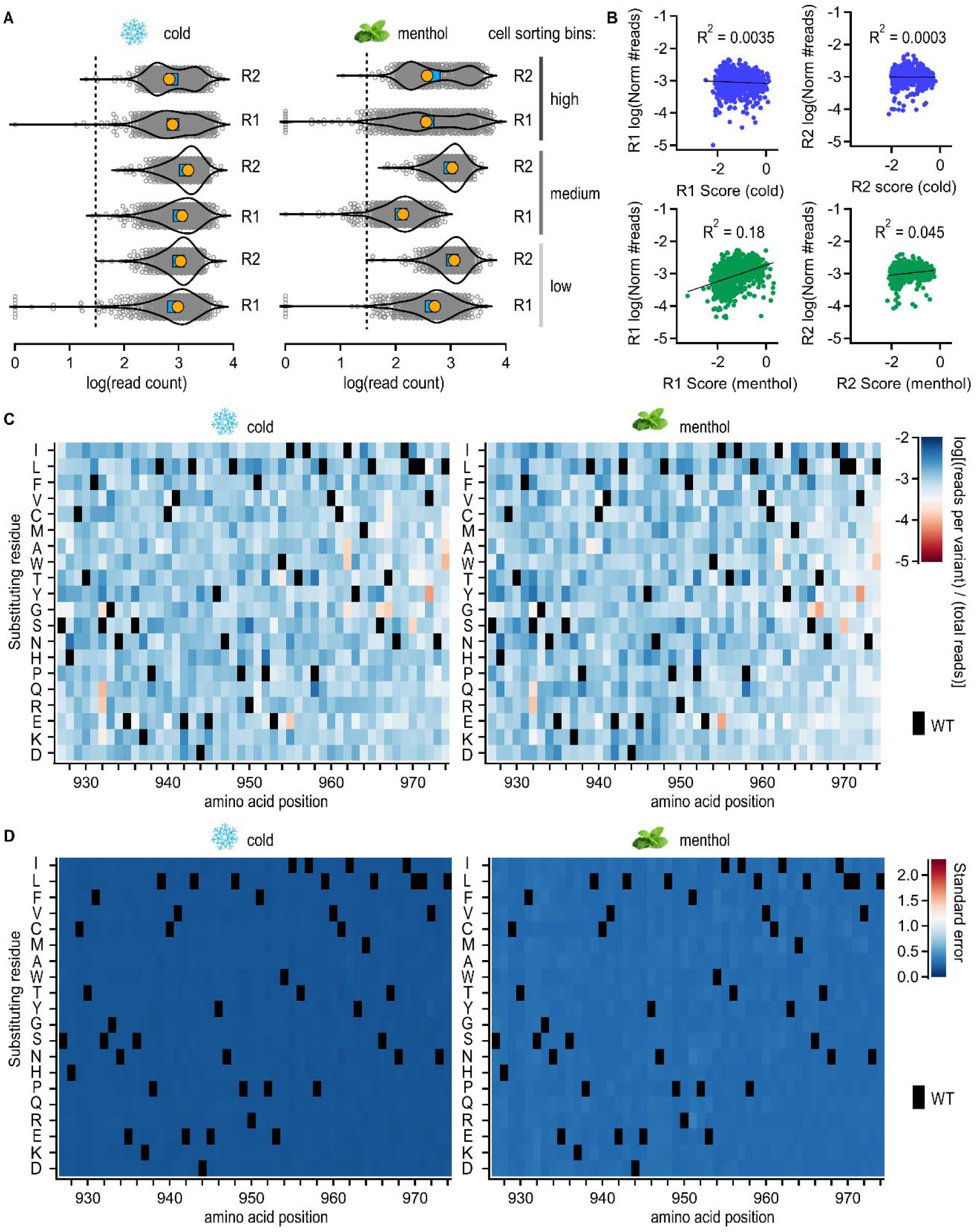
Read count distributions in sorted library cells. **(A)** Violin plots of the NGS variant read count distributions obtained from each of the groups of sorted cells in the two replicates (R1 or R2) and under the two experimental conditions (cold and menthol). Dashed lines denote a lower limit of 30 reads per variant. The mean and median of the distributions are shown in blue and yellow, respectively, and individual data points in gray. **(B)** Correlation between individual variant scores and the number of reads per variant relative to the rest of the variants summed across the three cell-sorting groups. **(C)** Heatmap of the logarithm of the number of total reads per variant across the two replicates relative to reads for the rest of the variants. Black squares denote the WT residues. **(D)** Standard error per variant as output by Lilace^53^.

**Extended Data Figure 3.**
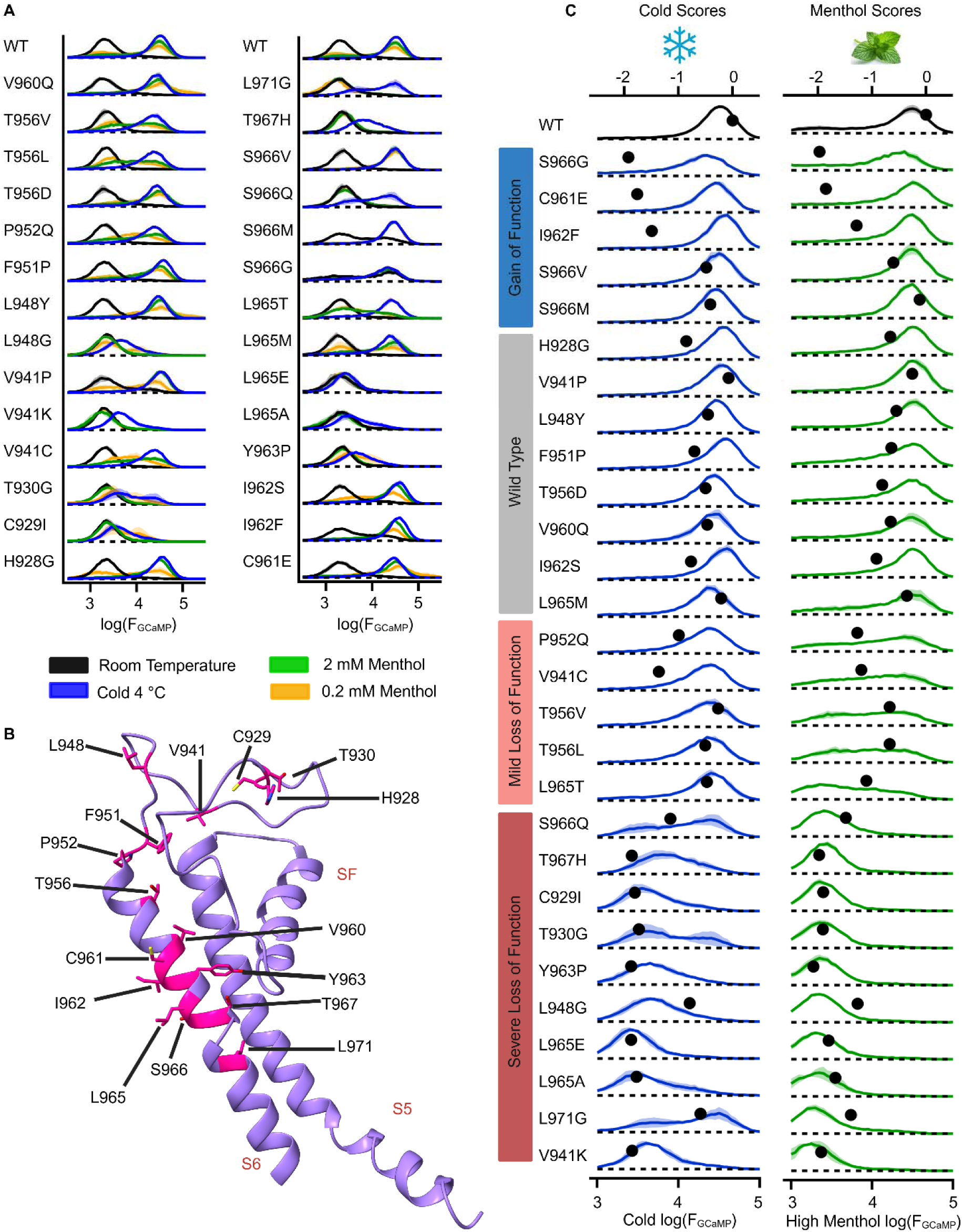
Benchmarking of the deep mutational scanning scores. **(A)** GCaMP6s fluorescence intensity histograms of mCherry^+^ cells expressing WT or mutant channels measured by flow cytometry at room temperature (black), or after simulation with cold (blue), 0.2 mM menthol (yellow), or 2 mM menthol (green). Data shown as mean ± SEM (n = 3). **(B)** Atomic model of the pore region of the TRPM8 D structure^15^ (PDB: 9B6D), showing each of the mutations in (A) in magenta. **(C)** Comparison of the WT and mutant GCaMP6s fluorescence intensity histograms for cold and 2 mM menthol (x-axis, bottom) and the deep mutational scanning scores of those same mutants (x-axis, top; black circles). Variants are grouped according to their phenotype as indicated by the color bars on the left side.

**Extended Data Figure 4.**
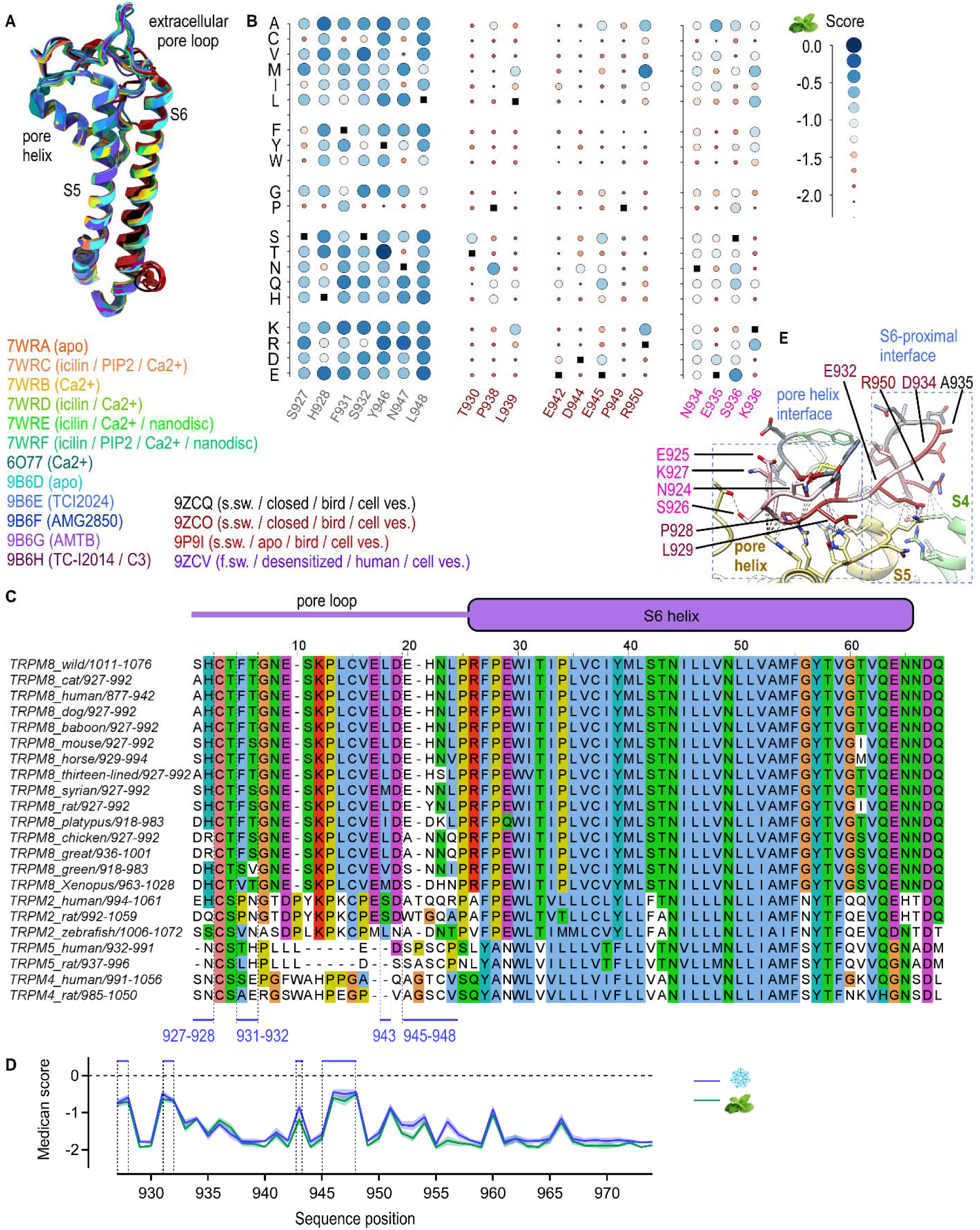
Sequence and structural conservation of the TRPM8 channel. **(A)** Structural alignment of the TRPM8 pore regions from 16 distinct structures as indicated by the colors and PDB labels. F.sw – fully swapped; s.sw – semi swapped. Only one subunit is shown. **(B)** Bubble plot of individual variant menthol scores for the same positions as in Fig. 3D. Scores are represented both by the radius of the circles and their color as denoted by the scale on the left on the heatmaps on Fig. 2A. Black squares denote WT residues. **(C)** Amino acid sequence alignment around the region from 927 to 974 in the rat TRPM8, including sequences from multiple TRPM8 orthologues from humans to amphibians as well as human and rat sequences for TRPM2, TRPM4, and TRPM5. Dashed lines and blue bars at the bottom denote regions where the TRPM8 sequences are not fully conserved. **(D)** Median deep mutational scanning scores per position, highlighting each of the regions with lower sequence conservation. The shading is the SEM of the scores across the 19 variants per position. **(E)** Structural model of an avian TRPM8 in the semi swapped closed state^31^ (PDB: 9ZCQ). Pore loop residues 927-951 are colored based on whether each position has a low (dark red), intermediate (pink), or high tolerance (gray) of substitutions based on the deep mutational scanning scores. The S4 helix of the same subunit as the pore loop is shown in green, and the pore helix and ion-selectivity filter (S.F.) of the adjacent subunit shown in sand. Dashed blue rectangles denote the pore helix and S6-proximal interaction interfaces between the pore helix and the rest of the protein. Dashed black lines denote non-covalent interactions between the pore loop and the rest of the protein.

**Extended Data Figure 5.**
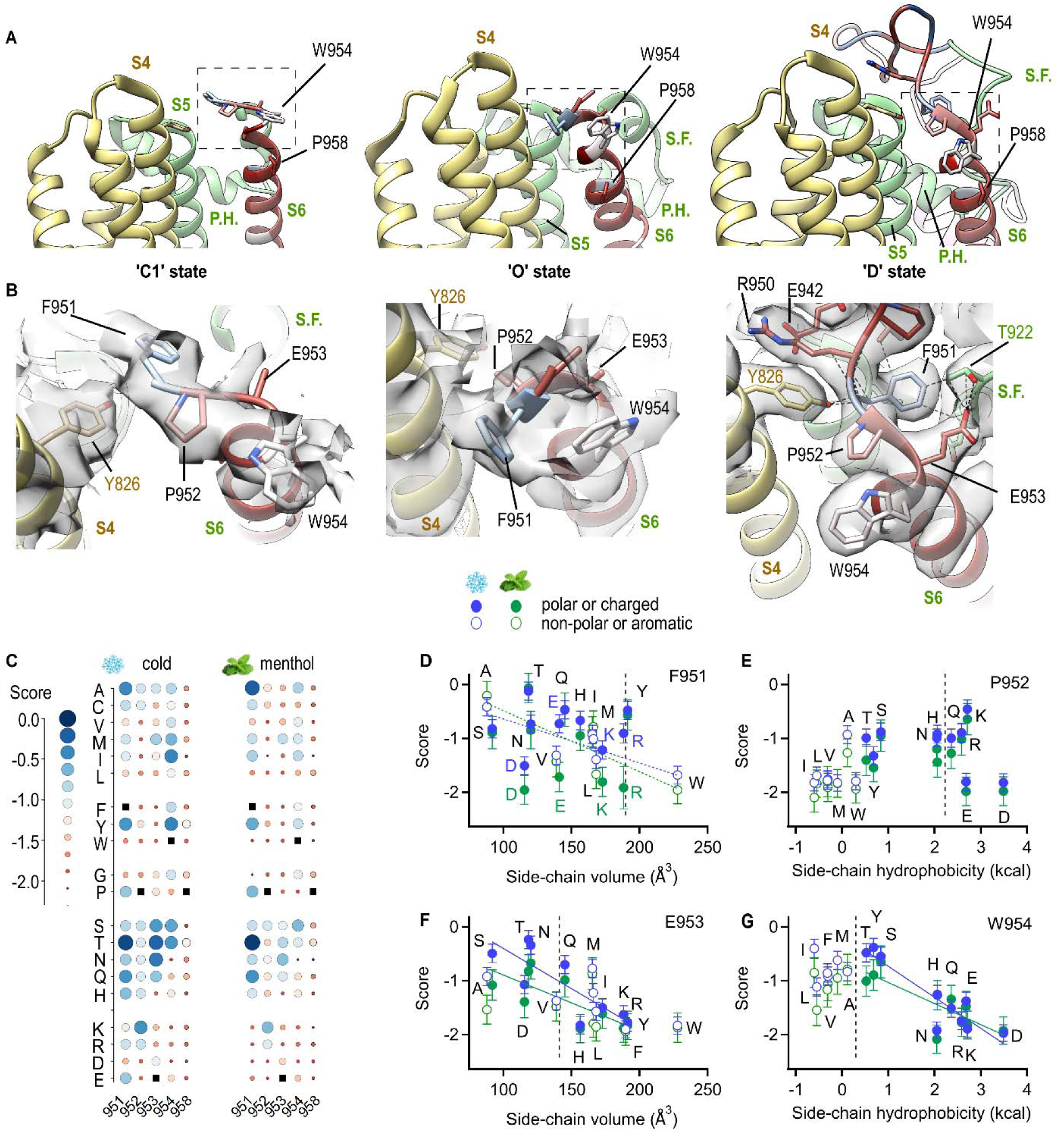
Gating-associated movements in the N-terminal end of the S6 helices. **(A)** Side view of the upper half of a portion of the transmembrane domain from C1^30^ (PDB: 8E4N), O^30^ (PDB: 8E4L), and D^15^ (PDB: 9B6D) structures of TRPM8. S6 residues of one subunit are colored by their median menthol score on a scale as in Fig. 2A, and the pore helix (P.H.) and selectivity filter of that same subunit are shown in green. S1-S4 helices of an adjacent subunit are shown in sand color. Dashed rectangles denote the approximate regions displayed in (B). **(B)** Magnified views of the structural models in (A), including the EM maps (C1, EMD-27893; O, EMD-27891; D, EMD-44255). Dashed black lines denote non-covalent interactions between the S6 and the rest of the protein. **(C)** Bubble plot of individual variant cold and menthol scores of the four N-terminal residues in the S6 as well as P958. Black squares denote WT residues. **(D-G)** Individual variant cold (blue) and menthol (green) scores as a function of substituting side chain volume or hydrophobicity^77^, distinguishing between polar and charged substituting amino acids (closed symbols) or non-polar and aromatics (open symbols). Error bars are the Lilace standard error per variant. Substitutions with proline and glycine were omitted. Vertical dashed lines denote the WT-residue properties.

**Extended Data Figure 6.**
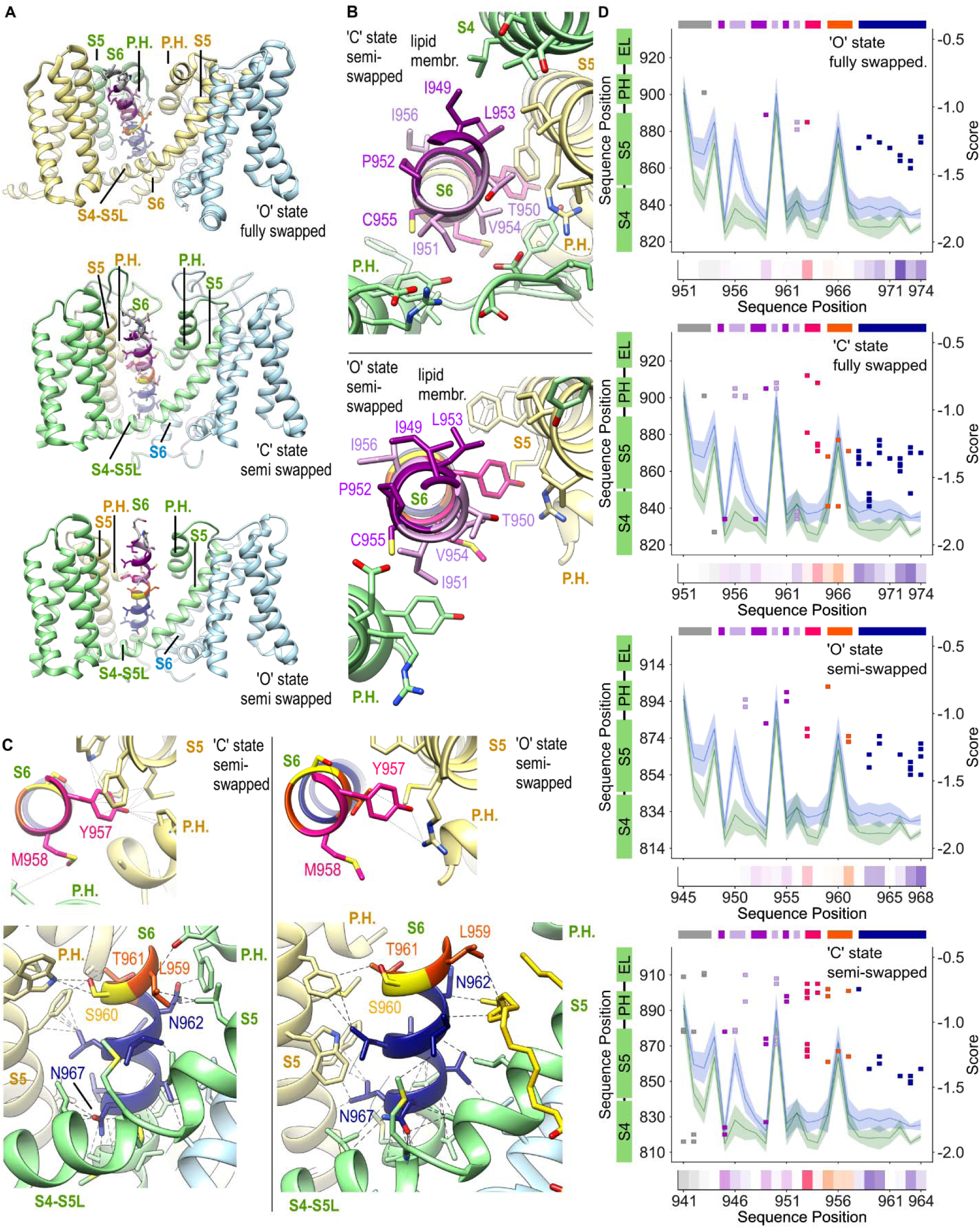
Structural determinants of gating across the S6 helices in distinct TRPM8 channel structures. **(A)** Transmembrane domain of TRPM8 in the fully swapped (f.sw) O^30^ (top, PDB: 8E4L) state, as well as in semi swapped (s.sw) C (middle; PDB: 9ZCQ) and O (bottom; PDB: 9PAR) states^31^ of an avian TRPM8. Subunit coloring is the same in all four complexes. **(B)** Depiction of an S6 helix and adjacent pore elements viewed from above in semi swapped (s.sw) C (PDB: 9ZCQ) and O (PDB: 9PAR) state structures of TRPM8^31^. S6 residues I955 to S966 are colored by helix region as in Fig. 4, the S5, pore helix (P.H.) and selectivity filter (S.F.) of the adjacent subunit colored in sand color, and the S4 and pore helices of the same subunit as the depicted S6 helix shown in green. **(C, top)** Structural depiction of the Y963 and M964 interaction interface in the two semi swapped structures with colors as in (A). **(C, bottom)** Depiction of an S6 helix and surrounding pore elements viewed from the membrane side in semi swapped structures of TRPM8. Residues and ribbons are colored as in (A). Dashed black lines denote non-covalent interactions between the S6 and the rest of the protein. **(D)** Map of residue-residue contacts between S6 helix residues 951-974 (x-axis) and the rest of the protein (y-axis, left) in a fully swapped open state of the human TRPM8 (top; PDB: 9PB5), a fully swapped closed state of the human TRPM8 (middle top; PDB: 9P8Y), a semi swapped open state of an avian TRPM8 (middle bottom; PDB: 9PAR), and a semi swapped closed state of an avian TRPM8 (top; PDB: 9ZCQ)^31^. Each color square on the map represents a contact and the colors denote each of the S6 regions as discussed in the text. The heatmap on the bottom insert denotes the number of total atomic contacts by the residues in the S6. The median cold and menthol scores per position are superposed (y-axis, right). The shading is the SEM of the scores across the 19 variants per position.

**Extended Data Figure 7.**
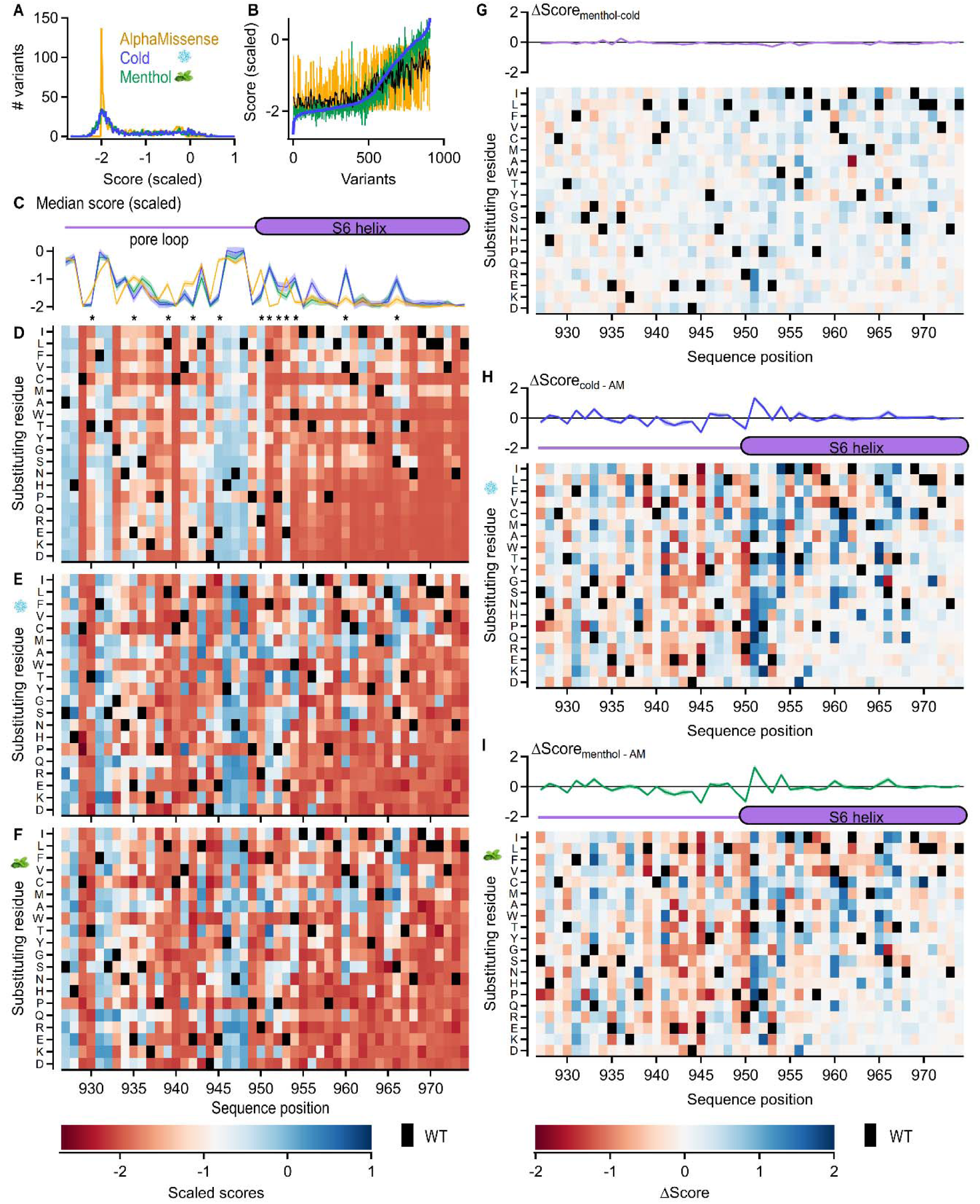
Comparison between experimental and predicted variant scores. **(A)** Predicted and experimental score distribution histograms for positions 927-974 after scaling to allow quantitative comparisons. AlphaMissense scores were scaled by a factor of -2. Experimental scores were scaled such that the peaks for the two components of the distribution were centered at -2 and 0. **(B)** Predicted and experimental score distributions after ranking all individual variants by their experimental cold scores from lowest to highest. The raw predicted scores distribution (yellow) and a smoothened predicted score distribution (black) are included. **(C)** Comparison of median predicted and experimental scores per position. The shading is the SEM of the scores across the 19 variants per position. Asterisks denote positions where the experimental and predicted scores diverge the most. **(D-F)** Heatmap of the scaled individual variant scores predicted by AlphaMissense^57^ (D) or obtained from cells stimulated with cold (E) or 2 mM menthol (F). Black squares denote WT residues. **(G-I)** Heatmaps of the difference scores between individual variant scores obtained under the two experimental conditions (G), or between the predicted and each of the experimental sets (H, I). The inserts at the top of each heatmap show the median difference score per position, and the shading is the SEM across the difference scores for the 19 variants per position. Black squares denote WT residues.

**Extended Data Figure 8.**
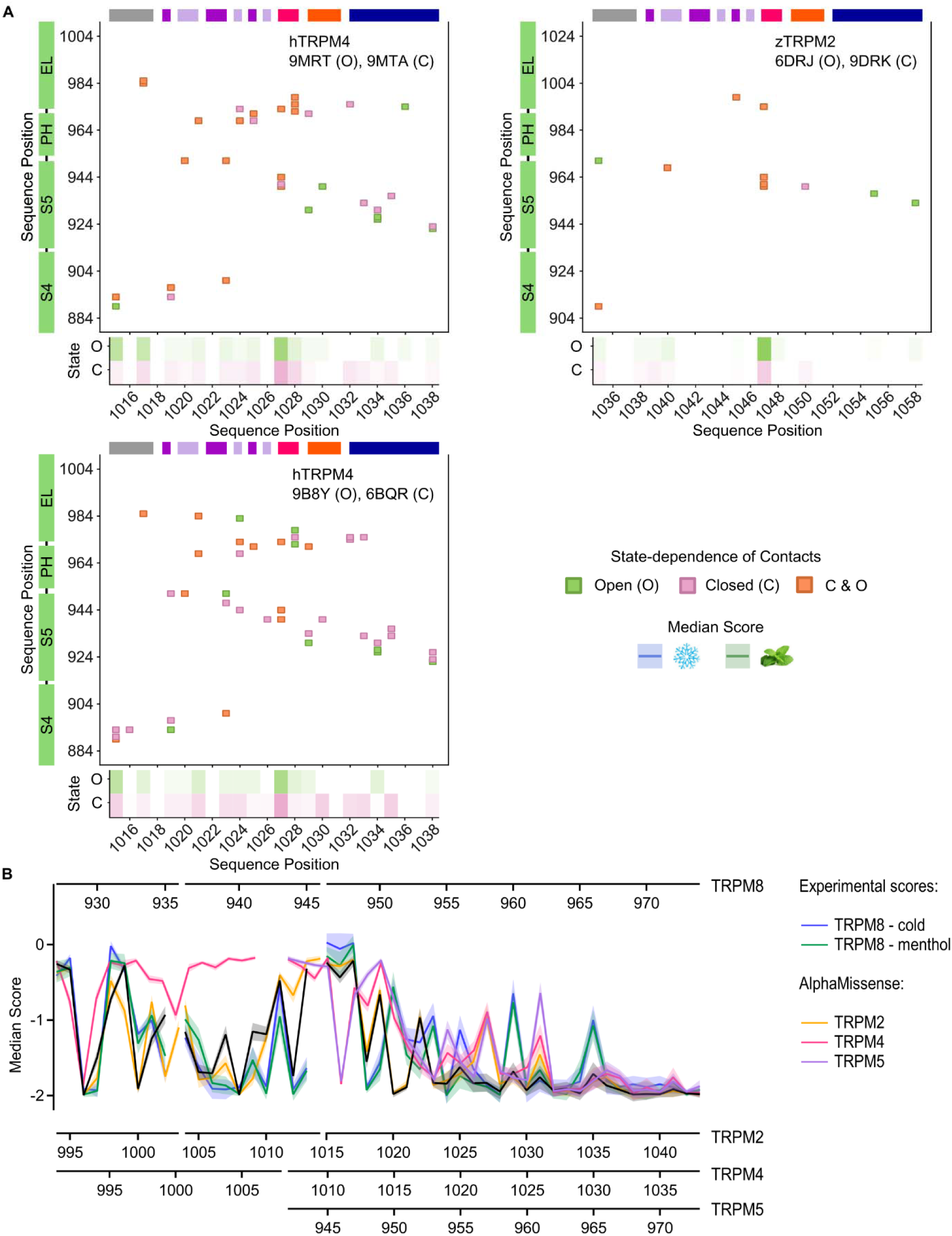
Structural determinants of gating in the S6 helix of other TRPM channels. **(A)** Map of state-dependent and independent residue-residue contacts involving the upper portion of the S6 helix and the rest of the protein, identified in closed and open-state structures of the TRPM2 (PDB: 6DRJ, 6DRK^68^) and TRPM4 (PDB: 9MRT, 9MTA^69^; 9B8Y^78^, 6BQR^79^) channels. Each color square on the map represents one residue-residue contact and the color of the square denotes its state dependence. The heatmap on the bottom insert denotes the number of total atomic contacts between each residue in the S6 and other residues in the rest of the protein. The portion of S6 that was included in this analysis was determined based on the amino acid sequence alignment between TRPM8 and the other channels (Extended Data Fig. 4B). (B) Median variant pathogenicity score obtained from scaled AlphaMissense predictions for all variants per position for the human TRPM2 (yellow), TRPM4 (magenta), and TRPM5 channels (purple), compared with the scaled predictions for TRPM8 (black) as well as the experimental scaled scores for cold (blue) and menthol (green). The shading is the SEM across the difference scores for the 19 variants per position.

**Extended Data Table 1.** Total number of sorted cells and variant read counts per sample and experiment. C_ **–** cold experiments; M_ – menthol experiments. _L – low GCaMP6s fluorescence group; _M – medium GCaMP6s fluorescence group; _H – high GCaMP6s fluorescence group.

| **Sorted Cell Count** |  |  |  |  |  |  |
| --- | --- | --- | --- | --- | --- | --- |
| **Replicate** | **C_L** | **C_M** | **C_H** | **M_L** | **M_M** | **M_H** |
| 1 | 136,381 | 503,182 | 413,916 | 500,000 | 500,000 | 382,529 |
| 2 | 152,170 | 500,000 | 500,000 | 500,000 | 500,000 | 500,000 |

| **Variant Read Counts** |  |  |  |  |  |  |
| --- | --- | --- | --- | --- | --- | --- |
| **Replicate** | **C_L** | **C_M** | **C_H** | **M_L** | **M_M** | **M_H** |
| 1 | 1,061,267 | 1,226,155 | 1,071,691 | 579,930 | 151,029 | 872,539 |
| 2 | 1,069,535 | 1,422,016 | 981,113 | 1,125,156 | 1,004,891 | 864,633 |

## Notes

### Competing Interest Statement

The authors have declared no competing interest.

